# Deep learning for mass extinction detection on fossilized phylogenies: power, limitations, and lessons for simulation-based phylodynamic inference

**DOI:** 10.1101/2025.10.20.683352

**Authors:** Minghao Du, Wenhui Wang, Jingqiang Tan, Joëlle Barido-Sottani

## Abstract

Detecting mass extinction events from phylogenies is a fundamental yet challenging task. While traditional likelihood-based methods are available, deep learning offers a powerful, simulation-based alternative. Here, we evaluate a deep learning approach using a novel hybrid model that combines Graph Neural Networks with Long Short-Term Memory networks. This model analyzes phylogenies—containing both extant species and fossils—simulated under a complex skyline Fossilized Birth-Death model that incorporates mass extinctions and fluctuating background rates. We validate the architecture’s effectiveness through ablation studies. Our investigation revealed that the stochasticity of the simulation was a primary obstacle, creating significant “label noise” that initially limited performance. A direct comparison showed our deep learning approach performed slightly better than Bayesian methods. It is robust to uncertainty in phylogenetic branch lengths and topology and generalizes to larger trees, but its performance degrades under model mismatch with higher background extinction rates. However, our work highlights a critical limitation: the model is highly specific to the definition of mass extinction it was trained on. Consequently, any modification to this definition necessitates retraining a new model from scratch. We conclude by summarizing the challenges and lessons learned for simulation-based inference in phylodynamics.

## 1 Introduction

The rapid advancements in artificial intelligence, particularly machine learning and deep learning [1], have led to significant breakthroughs across various scientific disciplines, establishing “AI for science” as a prominent research area [2]. Within phylogenetics, the application of machine learning is increasingly widespread [3,4]. A key development in this field is the use of machine learning for simulation-based inference to estimate parameters related to diversification rates in phylodynamic models, offering a powerful alternative to traditional likelihood-based approaches [5]. This methodology has been successfully applied across various phylodynamic models, accommodating both non-ultrametric and ultrametric trees [6–8]. The diverse ways of encoding phylogenetic trees—from summary statistics and specialized one-to-one mappings to treating trees as graph structures—have, in turn, led to the application of a wide array of neural network architectures, including multi-layer perceptrons, convolutional neural networks (CNNs), recurrent neural networks, and graph neural networks (GNNs) [9–11]. These approaches have found utility in various fields, such as epidemiology [12] and macroevolutionary studies [8]. While these approaches have shown great promise, applying them across the full spectrum of available phylodynamic models remains an active area of research.

Mass extinction events are extensively documented in the fossil record [13] and phylodynamic models have been developed to incorporate these major historical occurrences [14–16]. However, the very definition of a mass extinction is a subject of considerable debate, particularly regarding the magnitude of biodiversity loss and the timescale over which it must occur to qualify [17]. Geological history reveals a spectrum of patterns, from rapid, catastrophic crises like the Cretaceous-Palaeogene event to more protracted extinctions like the Late Devonian, which may have lasted for millions of years [17,18]. In this study, we focus on rapid, catastrophic events with a duration of less than one million years. Two primary approaches exist for modeling mass extinctions using phylogenetic trees: one involves a sudden, fixed survival probability occurring at a specific point in time, while the other employs a piecewise constant model that increases the extinction rate over a short duration [19]. Both methods can effectively be applied to fossil data through the Fossilized Birth-Death (FBD) model, a diversification model accounting for speciation, extinction and fossil sampling [20–22]. Despite the strong signature that mass extinctions can produce, their detection remains a persistent challenge [23,24]. This difficulty is exacerbated in more realistic models that allow background extinction rates to fluctuate continuously over time. Inferring mass extinctions under these complex conditions is substantially more difficult than under simpler models that assume constant background rates [16,22]. Indeed, recent studies using deep learning to classify diversification scenarios have shown that the phylogenetic patterns produced by mass extinctions are frequently confounded with those generated by high background extinction rates [25].

In this study, we investigate the detection of mass extinction events using deep learning within a simulation-based inference framework. We generate simulated phylogenetic trees under a skyline FBD model that incorporates mass extinctions and allows for fluctuating background extinction rates through time. Specifically, we model mass extinctions using a piecewise-constant approach, where the extinction rate is significantly elevated over a brief interval. Our work explores the performance of deep learning when applied to this highly flexible and realistic modeling framework. Our primary objective is to detect mass extinction events in phylogenetic trees. To achieve this, we developed a neural network architecture that couples Graph Attention Networks (GATs) with Long Short-Term Memory (LSTM) networks, chosen specifically for the features of the skyline FBD model. We then validated this design through ablation studies, systematically removing or replacing key architectural components like our custom time bin pooling mechanism and the use of virtual nodes to understand their impact on performance. We conduct the comparison of the performance of deep learning approaches against traditional Bayesian methods. Furthermore, we evaluate various factors that may influence model performance, such as tree size and the timing of mass extinction events. Recognizing that empirical phylogenies are reconstructions and thus inherently uncertain, we also systematically assess the model’s robustness to perturbations in internal node times (branch lengths) and topology. We also evaluate its behavior under conditions of model mismatch, specifically by using trees larger than those in the training data and higher background extinction rates. Although we initially viewed this as a simple binary classification problem, we encountered numerous challenges and unsuccessful attempts. Therefore, this paper aims to share our experiences and the lessons we learned, providing valuable insights to assist others in effectively applying deep learning for simulation-based inference in phylodynamics.

## 2 Data and Objectives

### 2.1 Data Simulation

To generate the data for our study, we employed a skyline FBD model. The FBD model is a stochastic process that jointly models speciation (birth), extinction (death), and fossil sampling, producing phylogenies that integrate both extant species and fossil specimens [20]. Crucially, these fossils can be incorporated as terminal tips or as sampled ancestors along branches, providing a richer depiction of evolutionary history. To create a challenging and realistic scenario, we modeled background diversification rates using a flexible horseshoe Markov random field (hsMRF) prior [26]. This locally adaptive prior models rates as being correlated between adjacent time intervals, which allows for smooth background fluctuations while retaining the flexibility to accommodate occasional rapid shifts.

We modeled mass extinctions as intervals of intense crisis by significantly elevating the extinction rate within a randomly selected 1-million-year time bin. This approach is methodologically significant as it ensures the entire process remains within the skyline FBD framework, whose parameters are theoretically identifiable [27]. To generate a more pronounced signal, we maintained low background speciation and extinction rates in contrast to the high rates during mass extinction events. Events were simulated probabilistically to ensure that approximately half of the trees experienced a mass extinction.

Using this strategy, we generated three large datasets for training, validation, and testing using the ReMASTER package [28] in BEAST2 [29]. For a comprehensive description of all simulation parameters, please see Section S1.

### 2.2 Objectives

We treat the detection of mass extinctions as a binary classification problem. The input for our model is a phylogenetic tree, and the output is a binary classification indicating the presence (1) or absence (0) of a mass extinction event within the phylogeny.

### 2.3 Model Architecture

A neural network framework is fundamentally composed of data and a model, and the architectural design of the model is a pivotal determinant of its performance. In this study, we adopted a hybrid model architecture that integrates GANs (a type of GNN) and LSTM networks (Figure 1). This approach is well-suited to our data, as GNNs are adept at processing graph-structured data like phylogenies, while LSTMs are specialized for handling sequential, time-series data. The resulting GAT-LSTM model represents a departure from the prevalent use of CNNs, which rely on grid-like data structures, in many prior phylogenetic studies (e.g., [6–8,10]). We chose this different path for two primary reasons. Firstly, given the intricate nature of our analytical tasks, we did not anticipate that standard CNNs would yield optimal performance. Secondly, applying CNNs to FBD trees would require the development of novel and potentially complex encoding methodologies to convert the tree structure into a grid-like format.

**Figure 1.**
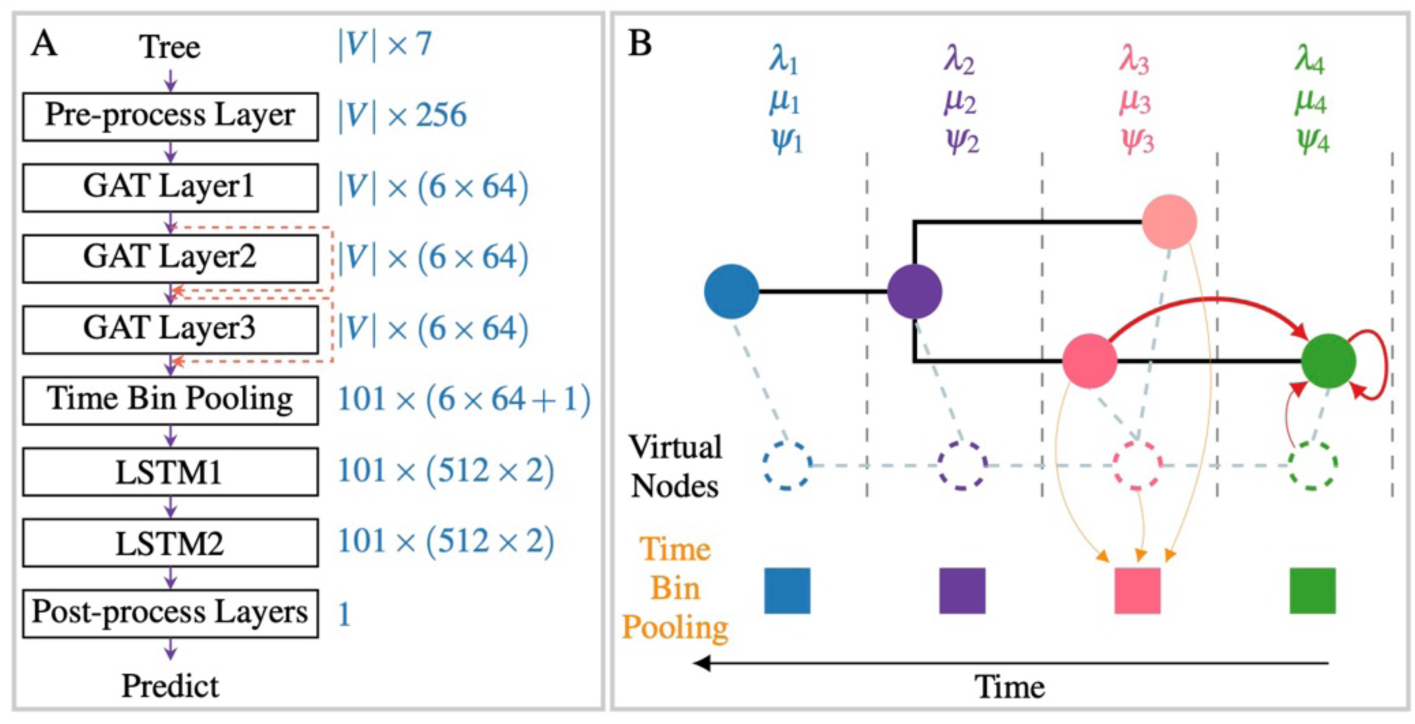
The architecture of the GAT-LSTM model. (A) The overall architecture of the model. Numbers indicate the feature dimensions, and |*V*| represents the total number of nodes in the tree (including tips and internal nodes). Red lines denote skip connections. For specific model parameters, see Table S2. (B) An illustration of the GAT-LSTM model applied to a simple skyline FBD tree, showing the incorporation of virtual nodes (dashed blue lines, see Section 2.3.3 for details) and the principle of time bin pooling. The red lines illustrate the information update for a single node within a GAT layer, where the line thickness corresponds to the attention weight assigned to different neighboring nodes (see Figure S2B and Section S2.2 for details). The orange lines represent the time bin pooling process for a single time bin (see Figure S2C and Section 2.3.4 for details).

#### 2.3.1 Graph Neural Network (GNN)

Because trees are a specialized form of graph, phylogenetic trees can be naturally conceptualized as graph-structured data. This makes them inherently suitable for GNNs and obviates the necessity for manual encoding procedures. Graph neural networks have seen extensive application across diverse scientific domains, encompassing a multitude of architectural frameworks (see [30,31] for a comprehensive review). Previous investigations have employed GNNs for diversification rate inference, wherein rudimentary GNN architectures demonstrated suboptimal performance relative to CNNs on elementary tasks [9]. However, subsequent research showed that through strategic modifications to the pooling mechanisms of GNNs—specifically tailored to accommodate the structural characteristics of phylogenetic trees and the requisites of the analytical tasks—their performance capabilities could be substantially enhanced [11].

GNN architectures typically comprise three main stages: pre-process layers (Section S2.1), GNN layers (Section S2.2), and post-process layers (Section S2.3). For a detailed exposition on the design space of GNN models, the reader is referred to [32]. Although we conducted preliminary evaluations for this project (Section S3), the identification of an optimal GNN architecture remains an area necessitating further in-depth investigation. This challenge arises from the vast design space of GNNs, where numerous configurations of layer types, pooling mechanisms, and network depth must be considered, making a systematic search for the best-performing model computationally intensive.

#### 2.3.2 Node Features

The first step in applying a GNN is to define the features for each node in the graph. The choice of which features to include represents a key modeling decision. In theory, a sufficiently powerful GNN could learn complex relational properties from a minimal set of inputs, such as branch lengths alone [31]. However, this approach increases the training cost, so we opted instead for a balanced approach by providing a set of core, informative attributes available within the phylogenetic tree. Each node in the tree was characterized by the following:

- Time before present (Ma).
- Associated time bin, corresponding to the same 1-Myr intervals used to define rate changes in the simulation (extant species are assigned to bin 0, with subsequent incremental assignment).
- Is an internal node (1 if yes, 0 if no).
- Is a terminal node (1 if yes, 0 if no).
- Is a fossil node (1 if yes, 0 if no).
- Is an extant species (1 if yes, 0 if no).
- Is a sampled ancestor fossil (1 if yes, 0 if no).
- Is a non-sampled ancestor fossil (1 if yes, 0 if no).
- Branch length to parent node (Myr).

While we provided this set of fundamental attributes, we did not augment the model with additional derivative features, such as those pertaining to a node’s child nodes, an approach taken by [10]. The optimal number of features for such tasks remains an open question. For instance, [10] showed that providing more information to a CNN can substantially improve model performance when training data is limited. Our approach therefore represents a middle ground between a minimalist input and a more heavily engineered feature set.

#### 2.3.3 Time Bin Virtual Nodes

In a standard GNN layer, information propagation is restricted to immediate neighbors. Consequently, enabling a node to receive information from more distant nodes requires an increase in the number of GNN layers. However, augmenting the layer count not only escalates computational demands but also introduces the risk of over-smoothing—a phenomenon where all nodes converge to nearly identical feature representations, thereby losing their discriminative capacity. This limitation is particularly pronounced in phylogenetic trees, which often exhibit large diameters; meaning the shortest path between the two most distant nodes in the graph is very long. To address this challenge, we incorporated a “virtual node” for each time bin to create efficient, long-range information pathways that reflect the core assumptions of our evolutionary model (Figure 1B). This architecture establishes two critical sets of connections: (1) Intra-Bin Connectivity: Each virtual node is connected to all real nodes within its corresponding time bin. This ensures these nodes can effectively exchange information, which is crucial for capturing the shared dynamics defined by the skyline model’s assumption that all species within the same interval share identical rates. This modification ensures all nodes in a bin are interconnected within a two-layer GNN structure. (2) Inter-Bin Connectivity: The virtual nodes are themselves interconnected sequentially with their neighbors. This emulates the temporal dependency of the Horseshoe Markov random field prior, which models rate correlations between adjacent time bins, reflecting a more realistic evolutionary dynamic. Together, these connections allow information to propagate rapidly across the entire tree—both within and between time periods— enabling nodes to receive information from distant parts of the phylogeny without requiring a deep GNN architecture. These virtual nodes were assigned temporal and time bin features consistent with their respective bins, while all other features were initialized to zero.

Intriguingly, our experimental findings indicated that the use of virtual nodes did not lead to a significant enhancement of model performance (Section S3 and Figure S3). A plausible explanation for this outcome is that even if substantial message propagation did not occur between nodes within the same time bin at the GNN layer stage, the time bin pooling mechanism, detailed subsequently, ultimately achieved the necessary aggregation of their information.

#### 2.3.4 Time Bin Pooling

Following the GNN layers, a pooling operation is required for graph-level prediction tasks, with the objective of aggregating information from all nodes into a single, unified representation (Figure 1B and Figure S2C). This process is conceptually analogous to the aggregation step within the GNN layers themselves. However, applying conventional global pooling methods— such as a simple mean, sum, or max pooling across all nodes—can be problematic. These approaches can lead to a significant loss of critical information pertinent to phylogenetic tree structures and their temporal dynamics, consequently diminishing neural network performance.

To address this challenge, we introduce a novel pooling strategy specifically designed for our skyline framework, termed “Time Bin Pooling”. The fundamental principle of this approach is to consolidate nodal information from the same time bin into a single vector representation. Specifically, for each time bin, we apply an average pooling operation to the features of all nodes that fall within that interval, thereby deriving a representative vector for that bin. This methodology effectively preserves the essential temporal information encapsulated within the phylogenetic time bins, concurrently mitigating the information loss often associated with traditional, non-structural pooling techniques. Indeed, when we compared our approach to a standard global mean pooling, we found that the latter yielded very poor results (Section S3 and Figure S3).

#### 2.3.5 Long Short-Term Memory (LSTM) Networks

Subsequent to the time bin pooling operation, rather than directly employing conventional post-processing layers (e.g., fully connected layers), we integrated Long Short-Term Memory (LSTM) networks. This decision was predicated on the inherent time-series nature of the information encapsulated within the time bins, as LSTMs are exceptionally proficient at capturing such temporal dependencies [33]. The input to the LSTM network comprises the sequence of feature vectors generated by the time bin pooling stage, and its output is a vector representation that encapsulates information across all time bins. This representation is subsequently amenable to downstream classification or regression tasks. We experimented with replacing the LSTM with two multilayer perceptrons (MLPs), but this led to a decrease in performance, likely because LSTMs are better able to capture the dependencies inherent in time-series data (Section S3 and Figure S3).

## 3 Experimental Evaluation

Armed with the simulated datasets and the developed model architecture, we embarked on an investigation to address our research objectives (see 2.2 Objectives). Throughout this endeavor, however, we encountered a series of significant challenges and methodological hurdles. While our efforts were ultimately successful, the challenges encountered were, in themselves, informative, providing a clearer understanding of the conditions under which our model performs effectively. Therefore, this section presents a comprehensive account of our experimental investigation. We will detail the findings and, equally importantly, discuss the valuable lessons derived from the attempts to resolve the objective. This analysis of both successes and obstacles aims to provide a transparent and holistic understanding of our model’s performance.

### 3.1 Preliminary Results

Our initial attempt yielded an accuracy of approximately 55% for classifying phylogenies both with and without mass extinction events. This result was underwhelming, as it represents only a marginal improvement over the 50% accuracy expected from random chance. Consequently, these initial findings were a source of considerable disappointment and prompted a deeper investigation into the potential causes of this suboptimal performance.

### 3.2 Refinement of Datasets

We initially hypothesized that the model’s suboptimal performance could be attributed to specific artifacts within the simulated datasets. The primary issue was the frequent weakness of the mass extinction signal, a problem stemming from two distinct sources: its temporal placement and its magnitude. Regarding placement, our initial constraint allowing extinctions after the first 20% of the tree’s history was insufficient, as events occurring too early were often difficult to detect. Regarding magnitude, the minimum added extinction rate of 0.4 was not always intense enough to be clearly distinguished from background rates.

Furthermore, the probabilistic nature of our simulation protocol introduced a source of noise. While designed to produce a single mass extinction, the process occasionally generated phylogenies with multiple such events. These rare but unintended outcomes introduced noise into the dataset, potentially confounding the model’s learning process.

To rectify these issues, we implemented a series of targeted refinements. To amplify the extinction signal, we simultaneously adjusted both its timing and magnitude. The event window was narrowed to occur between 30% and 80% of the tree’s origin time, and the intensity of the event was increased by setting the added extinction rate to a uniform distribution between 0.5 and 1.0 (Table S1). To eliminate the noise from multiple mass extinction phylogenies, we modified the simulation procedure. Instead of assigning a probability of mass extinction to each time interval, we now randomly select one specific interval for the mass extinction to occur (Table S1).

This revised protocol guarantees that each phylogeny in our dataset features exactly one, unambiguous mass extinction (Table S1 and Figure S1). Critically, it also ensures that our control datasets—those simulated without mass extinction—are genuinely free of such events, thereby removing ambiguous training examples with multiple extinctions and creating a more clearly defined classification task for the model.

Following this simulation strategy, we generated three distinct datasets. First, we simulated a comprehensive training dataset composed of 50,000 phylogenies with mass extinction events and 50,000 phylogenies without such events. We then generated a validation dataset and a test dataset to evaluate the model’s performance, creating an equivalent set for each. This resulted in 50,000 phylogenies with mass extinctions and 50,000 without for the validation dataset, and an identical set for the final test dataset.

After retraining the model with this refined dataset, we observed a marked improvement in its predictive power, with accuracy increasing to approximately 70% (Figure S4). However, while this represented a significant step forward, this level of accuracy was still not considered satisfactory for our purposes.

### 3.3 Tuning

With the dataset refined, we turned our attention to the model itself, systematically investigating whether performance was being constrained by the model. We conducted a broad range of experiments, including significantly increasing the model’s complexity (e.g., doubling key parameters, Section S4.1), performing extensive hyperparameter tuning (adjusting learning rates and dropout probabilities, Section S4.2), attempting to replace the core network architecture (using a Transformer block instead of LSTM, Section S4.3), and substantially increasing the volume of training data (from 100,000 to 300,000 phylogenies, Section S4.4).

However, none of these efforts yielded any significant improvement in performance; the model’s accuracy remained plateaued at around 70% (Figure S4). This result strongly indicated that the performance bottleneck was not due to a lack of model complexity, suboptimal hyperparameter settings, or insufficient data. We concluded that the root of the problem lay in a more fundamental aspect of our data or task definition.

### 3.4 Benchmarking Against a Bayesian Approach

To better contextualize our deep learning model’s performance, we benchmarked it against a traditional Bayesian method. We selected a representative subset of 100 phylogenies, each known to contain a mass extinction, and analyzed them using RevBayes [34]. To evaluate the Bayesian method under ideal prior conditions, it was configured with the same prior distributions used to simulate the training data (Section S5).

The results of this comparison were revealing. The Bayesian approach also struggled to reliably identify the extinction events. When using a lenient criterion for detection (2 log BF > 0), the method correctly identified an extinction in 74% of the phylogenies. However, this sensitivity came at the cost of precision, as it frequently inferred multiple, non-existent extinction events within the same tree. Conversely, applying a more stringent criterion (2 log BF > 2) reduced the overall success rate to just 61%.

This outcome provided a crucial insight: the task of identifying a single mass extinction from phylogenetic data is inherently difficult, even for established, powerful statistical methods. It suggested that our deep learning model’s performance was not as poor as we had initially thought. Rather, its ∼70% accuracy likely reflects the intrinsic difficulty of the problem itself. Given this insight, we decided not to proceed with an additional benchmark on 100 phylogenies that did not contain mass extinctions at this stage.

### 3.5 Identifying the Core Problem: Stochasticity and Data Artifacts

To diagnose the cause of the model’s performance plateau, we investigated the relationship between its accuracy and some key parameters, such as the timing (Figure S16) and extinction proportion of mass extinction events (Figure S17). This analysis revealed a fundamental issue: the stochastic nature of the simulation process was creating significant mismatches between the intended parameters and the realized patterns in the phylogenies. These data artifacts appeared in two forms.

First, the dataset intended to represent mass extinction contained phylogenies where no meaningful mass extinction had actually occurred. This discrepancy arose from two distinct sources. The first issue relates to low standing diversity, where a high extinction proportion does not translate to a high absolute number of extinctions. A high-magnitude extinction has little practical impact if it occurs when the total number of species is already small. For instance, a mass extinction event programmed with a high 80% extinction proportion affecting a phylogeny of only five species results in the loss of just four lineages. While proportionally significant, the absolute number of extinctions is small and may not produce the strong phylogenetic signal characteristic of a true mass extinction. The second, more subtle issue stems from the stochastic nature of the simulation process itself, where the realized extinction proportion can deviate significantly from the programmed parameter. Our simulation sets a probability of extinction for each lineage, not a fixed number that must go extinct. Consequently, random chance can lead to a realized outcome that is far less severe than intended, especially when the number of species is modest. For example, consider a simulation with 20 species where we set the extinction proportion to 50%. This is analogous to flipping 20 fair coins; while the expected outcome is 10 extinctions, it is statistically plausible to observe only two extinctions. In this case, the realized extinction proportion is just 10%. In both scenarios, the phylogeny was labeled as containing a mass extinction (based on the input parameters), but it lacked the corresponding distinct signal in the final data, fundamentally misleading the model during training.

Second, the control dataset, intended to be free of mass extinctions, contained phylogenies that misleadingly appeared to have them. While the maximum background extinction rate was capped at 20%, periods of low species richness could stochastically experience the loss of a high proportion of their lineages. This created a phylogenetic signature that was functionally indistinguishable from a genuine mass extinction event.

These stochastic discrepancies introduced significant label noise, or “dirty data”, into our training set. We concluded that this fundamental conflict between the intended simulation category and the actual resulting data was the primary factor limiting the deep learning model’s performance.

### 3.6 Redefining Mass Extinction

To rectify the issue of data-label mismatches, we developed a new, quantitatively robust definition for a mass extinction based on its realized impact within a phylogeny. Henceforth, we defined a mass extinction as an event where, within a single 1-million-year interval, two conditions were met: (1) at least 16 species went extinct, and (2) this loss represented more than 50% of the standing diversity at the beginning of that interval. The number 16 was chosen because, in the simulated data for trees without mass extinctions, one tree had 15 species go extinct, representing a 50% loss. Therefore, this choice ensures that in the training set, all trees simulated without a mass extinction are classified as such. We explored alternative thresholds for defining a mass extinction, specifically setting the minimum species loss at 11 and 21. Neither of these values yielded the same level of accuracy as the 16-species benchmark. The lower threshold of 11 species was overly inclusive, frequently misidentifying trees without a mass extinction as having experienced one. Conversely, the higher threshold of 21 species was too stringent, often failing to detect genuine mass extinction events. Consequently, the 16-species definition represents the most robust and balanced threshold for identifying mass extinctions in our analyses.

This definition, based on the actual outcome in the phylogeny rather than the intended simulation parameters, allowed us to re-curate our datasets with high fidelity. Applying this stringent filter revealed the profound extent of the original problem. Of the 250,000 phylogenies simulated without a programmed mass extinction, only a single tree met our new criteria, confirming the filter’s effectiveness at eliminating false positives. Conversely, of the 250,000 phylogenies simulated with an intended mass extinction, only 137,423 (approximately 55%) actually produced a phylogenetic pattern consistent with our revised, more realistic definition. This re-classification effectively purged the ambiguous and misleading examples from our training data.

### 3.7 Final Results

Following this rigorous data re-labeling, we retrained the model on the refined dataset. The new training set was imbalanced (82,655 phylogenies with mass extinctions and 217,345 without). To compensate for this, we implemented a weighting scheme during training that gave proportionally more importance to the underrepresented mass extinction cases. This time, the model’s accuracy improved significantly to approximately 80% (Figure 2), with comparable performance across the training, validation, and test sets (Figure S5).

**Figure 2.**
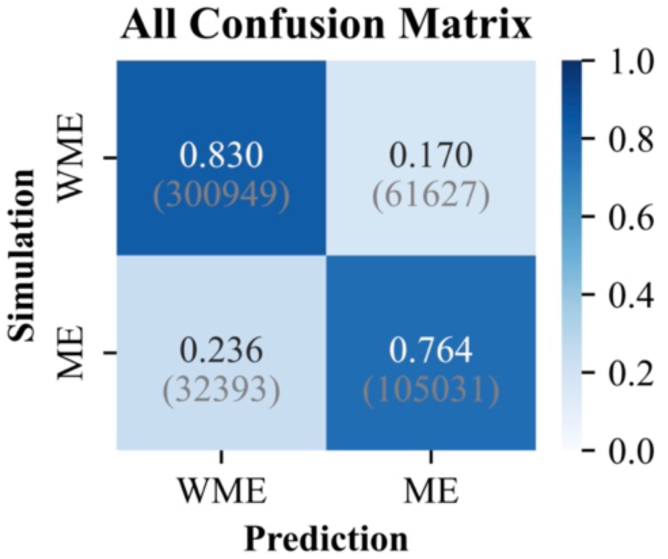
Confusion matrices illustrating the model’s accuracy on the training, validation, and test sets under the redefined criteria for mass extinction. ME: Mass Extinction; WME: Without Mass Extinction. Numbers indicate the proportion of predictions for each true class, with the absolute counts of phylogenies in parentheses.

#### 3.7.1 Bayesian Benchmark

Inspired by [7], we conducted a final comparison between our refined deep learning model and the Bayesian method. We first selected the most challenging cases: 100 phylogenies with mass extinctions that the deep learning model failed to detect (false negatives), and 100 phylogenies without such events that it misclassified (false positives). Given the limited dataset of 200 phylogenies, we did not directly compare accuracy. Instead, we compared the Bayesian method’s Bayes factor with what we term the deep learning score. The deep learning score, derived from a sigmoid function, ranges from 0 to 1 [35]. A score below 0.5 indicates the absence of a mass extinction event, while a score above 0.5 suggests its presence. Generally, a higher score reflects greater confidence from the deep learning model in detecting a mass extinction, and can be interpreted as the deep learning equivalent of a Bayes factor. The results revealed a moderate correlation between the two methods, but their outcomes were not consistent, with several notable discrepancies observed (Figure S6).

To better understand the respective failure modes of our deep learning model and the Bayesian method, we conducted a detailed post-mortem on four phylogenies where their conclusions diverged significantly. For each case, we compared their inferences against the true, simulated history of speciation, extinction, fossilization, and diversification (Figure S7–Figure S10).

In two of these cases, the deep learning model failed to detect a genuine mass extinction that the Bayesian analysis correctly identified. A closer look revealed that the number of species lost in these events was relatively low (19 and 18 species, respectively; Figure S7 and Figure S8), falling just below or near the threshold of our revised definition (16). This suggests the model learned to treat these borderline events as background noise. It is worth noting, however, that in one of these instances, the extinction detected by the Bayesian method did not correspond to the true simulated event time, highlighting its own potential for error (Figure S7).

One particularly informative case arose from a simulated evolutionary history that included a mass extinction event, eliminating 87 species, or 49.43% of the total biodiversity (Figure S9). Our deep learning model flagged this event with a very high mass extinction score. In direct contrast, a conventional Bayesian analysis did not detect a mass extinction. The Bayesian method’s conclusion was technically correct based on our working definition, which requires that a mass extinction must eliminate over 50% of species. The Bayesian approach performed as expected in this scenario. When the background extinction rate (estimated at approximately 0.1) is subtracted from the observed 49.43% extinction, the effective proportion of the extinction pulse is closer to 40%. For the Bayesian inference framework, this 40% extinction proportion is substantially lower than the required 50% threshold. Given the high species richness in the simulation, this distinction was pronounced enough for the method to correctly classify the event as a non-mass extinction under our strict definition. Curiously, however, the Bayesian analysis also failed to infer any significant increase in the background extinction rate for that period. This observation suggests that our definition of mass extinction may still be suboptimal, as it does not fully capture such significant, albeit borderline, events. This specific example highlights a critical challenge in applying machine learning to scientific problems: the success of the model is fundamentally tied to how precisely and accurately the learning objective—in this case, the very definition of a mass extinction—is defined.

Finally, we examined a phylogeny with no mass extinction that the model incorrectly flagged (Figure S10). This tree was characterized by a prolonged period of elevated extinction rates and continuously declining diversity beginning around 45 Ma, though the extinction proportion in any single time bin remained low. We infer that this pattern of a “slow-burn” crisis was underrepresented in our training data. As a result, the model, lacking a learned category for this specific dynamic, misclassified it as the most similar phenomenon it knew: an abrupt mass extinction.

Upon examining the phylogenies where deep learning predictions were entirely erroneous, we observed that while deep learning and Bayesian methods exhibited some correlation, this relationship was weaker than anticipated. We hypothesized that this was not a general feature of the methods, but rather an artifact of our sample selection. By focusing exclusively on the model’s failures, we were analyzing a collection of phylogenies with inherently ambiguous or challenging characteristics. To test this hypothesis and obtain a more representative comparison, we performed an additional benchmark analysis on a new, randomly selected set of 200 phylogenies (100 with and 100 without a mass extinction). As anticipated, the correlation between the deep learning and Bayesian inferences was substantially stronger in this unbiased sample. Nevertheless, a number of phylogenies still yielded conflicting predictions, prompting a closer examination of some specific cases (Figure 3A).

**Figure 3.**
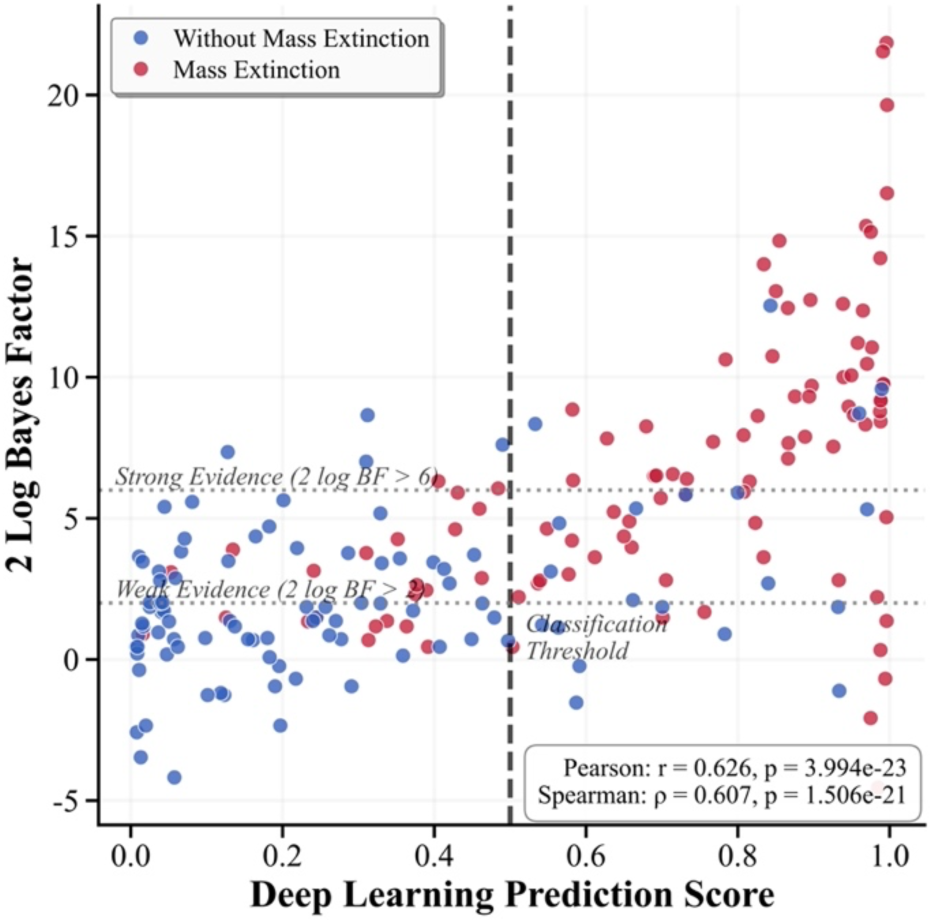
Comparative scatter plot illustrating the performance of Deep Learning versus Bayesian inference on the confidence of detecting mass extinction events. Colours indicate whether a mass extinction was present in the phylogeny.

Our analysis of these remaining disagreements revealed patterns consistent with our previous findings, further clarifying the distinct operational biases of each method. For instance, in cases where the extinction proportion was close to our 50% threshold but the absolute number of extinct species was large, the deep learning model consistently inferred a mass extinction while the Bayesian method did not (Figure S11–Figure S13). Conversely, when the extinction event was proportionally significant but involved a small absolute number of species, the deep learning model often failed to detect it (Figure S14). Finally, this new sample also contained a particularly informative anomaly: a phylogeny with no true mass extinction where our deep learning model correctly reported a negative result, but the Bayesian method erroneously inferred a significant extinction event (Figure S15). This highlights that both methods are susceptible to error, albeit on different types of phylogenetic patterns, and reinforces that neither approach is infallible.

#### 3.7.2 Factors Influencing the Accuracy of Mass Extinction Detection

To identify the factors that influence the model’s predictive power, we systematically analyzed the relationship between its accuracy and scores against key phylogenetic parameters. For the most part, the results aligned with our expectations. The model’s performance improved with a greater number of tips (Figure S18), more fossils (Figure S19), a larger absolute number of extinct species (Figure S22), and higher pre-extinction diversity (Figure S23). Timing was another critical factor: extinctions occurring later in a tree’s history were detected with much higher accuracy, whereas the model had very little predictive power for early events (Figure S24). We also observed some unexpected, complex relationships, such as a U-shaped curve between accuracy and the number of extant species, which may be an artifact of sampling bias in our simulation protocol (Figure S20). For a detailed breakdown of how these factors influence model performance, see Section S6.

## 4 Assessment of Robustness to Phylogenetic Uncertainty

Real-world phylogenetic analyses are subject to inherent uncertainty, and empirical datasets may not perfectly align with the distribution of the training data. It is therefore critical to evaluate the robustness and generalizability of our deep learning model. The computational speed of the trained model offers a significant advantage, enabling comprehensive and rigorous assessments that are often computationally prohibitive for traditional Bayesian methods.

### 4.1 Robustness to Phylogenetic Uncertainty

Phylogenetic trees are reconstructions, not ground truths, and are thus subject to uncertainty in both their branch lengths (internal node times) and topology. To assess our model’s performance in the face of this uncertainty, we systematically perturbed the internal node times and topologies of our test phylogenies, both individually and simultaneously (Section S7).

Our results show that the deep learning model is robust to this type of uncertainty. Although the model’s raw prediction score diverged slightly with increasing perturbation, the final classification—whether a mass extinction occurred or not—remained highly stable (Figure S25 and Figure S26). Even under the most extreme scenarios tested, the vast majority of perturbed phylogenies yielded the same prediction as the original, unperturbed tree (Figure 4 and Section S7.3). This stability suggests the model has not overfitted to specific features of the training data but has instead learned the broader phylogenetic patterns characteristic of mass extinctions.

**Figure 4.**
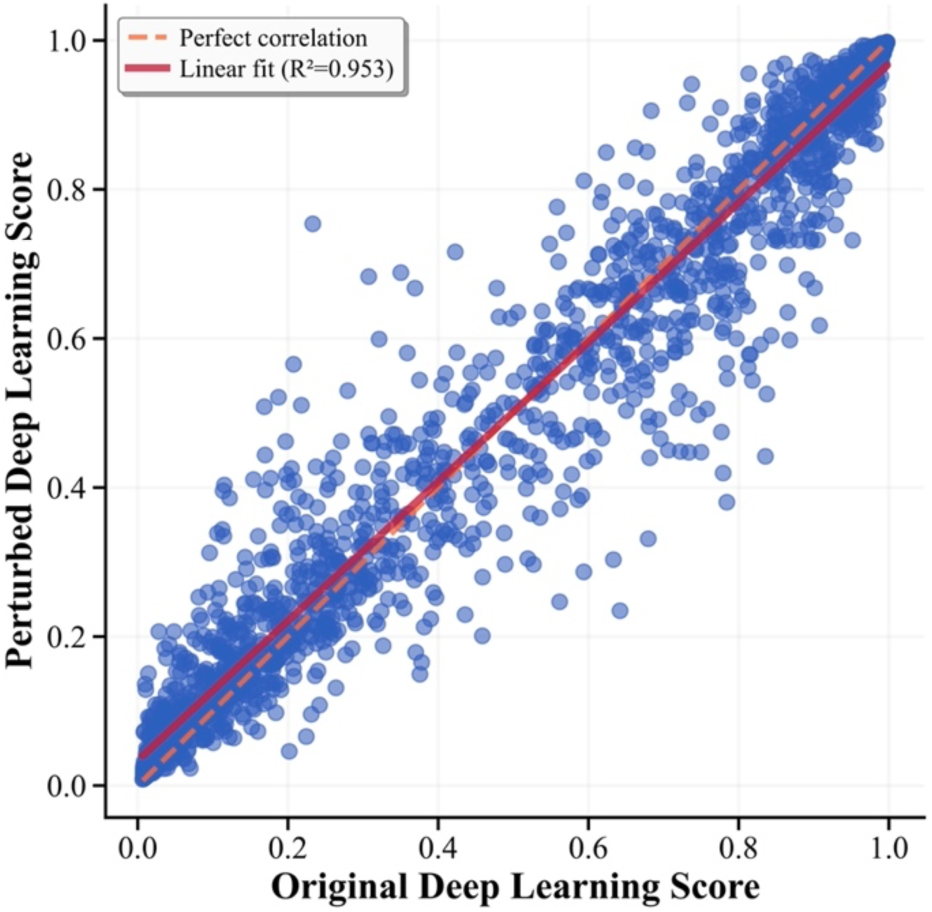
Assessing the robustness of the Deep Learning model’s predictions to phylogenetic uncertainty. The scatter plot compares the model’s estimates for the confidence of detecting mass extinctions, derived from original trees versus those from trees with simultaneously perturbed internal node times and topologies.

### 4.2 Performance Under Model Mismatch

We next evaluated the model’s ability to generalize to data drawn from outside its training distribution. We investigated two model mismatch scenarios: phylogenetic trees of a larger size and trees simulated under higher background diversification rates (Section S8).

When applied to larger trees (2,200–4,000 tips, compared to the 200–2,000 tip training range), the model’s accuracy in detecting the presence of a mass extinction improved (Figure S5K). We attribute this successful generalization to our network architecture; specifically, the time-bin pooling strategy and the local nature of the GNN layers allow the model to effectively handle an increased number of lineages within each time interval without altering its core computational structure.

In contrast, the model’s performance degraded substantially when tested on phylogenies with higher background extinction rates. Under these conditions, the model exhibited a high false-positive rate, misclassifying stochastic fluctuations in the background extinction rate as mass extinction events (Figure S5L). This result highlights a key limitation: the model’s predictive capabilities are highly dependent on its training distribution, as the very definition of a mass extinction becomes less robust when the distinction between catastrophic events and background noise is blurred.

## 5 Lessons Learned

### 5.1 Capabilities of Deep Learning

Our findings demonstrate that, under the conditions defined for our study, the deep learning model is a powerful tool for analyzing mass extinction events. It performs remarkably well in detecting the presence of a mass extinction, and a direct comparison reveals that its performance is comparable, and in some respects superior, to conventional Bayesian methods. This underscores a critical insight: when the predictive target is well-defined, deep learning can achieve high performance in complex phylogenetic analyses. However, while the inferences from deep learning and Bayesian methods show a moderate to strong correlation, their outcomes are not entirely consistent, with a number of phylogenies yielding conflicting predictions. These mismatches suggest that the two approaches may capture different features of the phylogenetic data; they are not mutually exclusive but rather complementary. Future research should focus on a deeper comparison of their similarities and differences to better understand the phylodynamic information contained within phylogenetic trees.

A key lesson from our study for simulation-based inference is the critical importance of inspecting simulated data to define a precise predictive target and eliminate “label noise”.

Due to the stochastic nature of the simulation process, unexpected outcomes are common; for instance, a simulation programmed to include a mass extinction may result in a phylogeny lacking any meaningful extinction signal, while a control simulation might stochastically produce a pattern resembling one. We found this “label noise” to be the primary factor limiting our model’s initial performance. Once we established a new, quantitatively robust definition for a mass extinction—based on the realized number of extinct species and the proportion of standing diversity lost—and re-labeled our data accordingly, the deep learning model’s accuracy improved dramatically. This ability to train a model on a custom, outcome-based predictive target is a significant advantage of the deep learning approach. While implementing such multi-faceted, user-defined criteria can be challenging within a conventional Bayesian framework, it is a straightforward process in deep learning that, as we have shown, can lead to substantial performance gains.

This same principle applies not only to classification but also to regression tasks. For example, if the goal was to predict the intensity of a mass extinction, training a model to predict the input extinction rate parameter would be misleading. A more accurate and robust approach would be to predict the realized outcomes, such as the actual percentage of species that went extinct in an interval or the absolute number of extinctions. This is because even a very high simulated extinction rate may not produce a strong signal if the standing diversity is low or if stochasticity results in fewer actual extinctions than expected. The rate parameter is stochastic in its effect, whereas the realized number of extinctions and the resulting biodiversity curve are the deterministic outcomes that are truly encoded in the phylogeny. This issue of simulation stochasticity should also be a concern in likelihood-based studies, as methods are often validated through simulation, and if the simulations themselves are problematic, the evaluation of the method will also be flawed [36].

The model’s impressive robustness to phylogenetic uncertainty and its unexpected scalability also stood out. Even when subjected to perturbations of branch lengths and topology, the model’s final predictions remained highly stable. This indicates that the deep learning model has genuinely learned the characteristic features of a mass extinction from the phylogenetic trees, rather than simply memorizing the training data, which makes its results highly credible. Moreover, when applied to trees larger than those in the training set, the model’s performance in detecting mass extinctions did not decrease but actually showed a slight improvement. This suggests that, with a properly designed neural network architecture, deep learning models can be scalable to a certain extent for this type of problem. How to design scalable network architectures for specific phylogenetic questions is therefore a promising area for future research.

Furthermore, the deep learning approach offers a profound practical advantage in its computational efficiency. Once a model has been trained, it can be applied to new data almost instantaneously. This speed is not merely a convenience; it unlocks the ability to conduct comprehensive and rigorous evaluations of the model’s behavior that are often computationally prohibitive for traditional Bayesian methods. For instance, it becomes feasible to systematically assess the model’s robustness to phylogenetic uncertainty, explore the factors that influence its performance, and test its scalability on datasets of varying sizes, thereby providing a much deeper understanding of the model’s strengths and limitations.

### 5.2 Limitations of Deep Learning

Despite its considerable strengths, the deep learning approach is not without significant limitations, and one of its key advantages—the ability to train on a custom, well-defined predictive target—is paradoxically also a critical weakness. While creating a precise, outcome-based definition for a mass extinction dramatically improved our model’s performance, such definitions are inherently subjective. For instance, one could question why the threshold was set to a loss of at least 16 species and not 20, or an extinction proportion of 50% rather than 60%. This subjectivity leads to a significant practical problem: there is no single, universal model for all definitions of mass extinction [17]. Any modification to the definition requires the entire workflow to be repeated—a new model must be trained from scratch and its robustness and performance factors must be completely re-evaluated. This reliance on a rigid, custom definition is also precisely why the model fails when confronted with data outside its training distribution. When we tested our model on phylogenies simulated with higher background extinction rates, the distinction between normal fluctuations and catastrophic events became blurred. Many stochastic background extinctions met the strict criteria of our custom definition and were consequently misclassified as mass extinctions, leading to a high false-positive rate and a substantial reduction in the model’s predictive capabilities.

Furthermore, creating a single, universal model applicable to the full range of empirical datasets presents a nearly impossible computational burden. The parameter space of complex phylodynamic models is vast, and a truly general model would need to be trained on simulations covering every plausible combination of scenarios, leading to a combinatorial explosion of possibilities. Our study, by necessity, introduced several constraints to make the problem tractable. For example, we limited background diversification rates to a narrow range (0–0.2), whereas real-world rates can be much higher, and assumed complete sampling of living species (an extant sampling probability of 1.0), while in practice sampling fractions can vary widely and be biased. Similarly, we restricted the tree origin time to a window between 80 and 100 million years and focused our final model on detecting a single mass extinction, yet empirical histories can span vastly different timescales and include multiple mass extinction events. Each of these parameters represents another dimension in an already high-dimensional space. To properly train a model to handle this immense complexity would require simulating billions of trees and developing a deep learning architecture with a corresponding scale of billions of parameters—a task well beyond current practical limits.

A final challenge lies in navigating the vast design space of the deep learning workflow itself. The development process involves a multitude of choices regarding model architecture, training metrics, and hyperparameters. This makes it difficult to gauge performance objectively, a problem compounded by deep learning’s general lack of interpretability. Unlike Bayesian methods, it can be challenging to determine precisely which phylogenetic features a model has learned, making it hard to diagnose the reasons for its failures. For example, our model’s initial 70% accuracy was first viewed as a failure. Only after benchmarking it against a traditional Bayesian method—which itself struggled to achieve a comparable success rate—did we understand that this performance reflected the intrinsic difficulty of the problem rather than a flaw in our model. Furthermore, it is incredibly difficult to predict which modifications will actually improve results. Our attempts to boost performance by doubling the model’s size, changing its core architecture, and tripling the training data all failed to yield any significant gains. This unpredictability highlights the need for a unified, standard dataset for simulation-based phylodynamic inference, which would allow for a more direct and rigorous comparison between different deep learning architectures.

### 5.3 (Perhaps) More Effective Applications of Deep Learning

Inferring diversification rates presents a unique challenge for deep learning. Unlike many applications where vast, pre-labeled datasets are readily available, phylogenetic analyses typically rely on simulation-based inference. In this paradigm, researchers must first construct a generative model—such as one of the many variants of the FBD model—and then use it to simulate the data needed for training. This workflow is immensely labor-intensive, as a new deep learning model must be trained for every distinct biological model a researcher wishes to test. As we have discussed, creating a single, universal deep learning model that is applicable to all empirical datasets is a computationally impossible goal.

A more effective and pragmatic application of deep learning in this domain may therefore lie not in replacing traditional methods, but in augmenting them. A hybrid approach that combines the speed of deep learning with the flexibility of MCMC methods is particularly promising. In such a framework, deep learning could be used to dramatically accelerate the parameter estimation step within an MCMC analysis. This would preserve the essential ability for experts to formulate and test novel biological hypotheses using custom models, while benefiting from rapid inference. Alternatively, following the example of [37], deep learning could be integrated into the model itself to account for non-linearities and interactions among variables.

Furthermore, by using deep learning as an accelerator rather than a standalone inference engine, it becomes far more feasible to train models on a case-by-case basis for smaller, specific datasets. This would make the power of deep learning more accessible and adaptable for targeted, hypothesis-driven research in systematics.

## Acknowledgements

We are grateful to Daniele Silvestro, Torsten Hauffe, Sophia Lambert, and Am’elie Leroy for their insightful discussions. We thank the Morlon team at IBENS and Rachel Warnock at FAU for their very helpful suggestions and feedback on an earlier draft of this manuscript.

## Funding

JBS received support from the European Union’s Horizon 2020 Research and Innovation Programme under the Marie Sklodowska-Curie grant agreement No. 101022928. MD received support from the China Scholarship Council (grant number 202306370127). JT received support from the National Natural Science Foundation of China (grant number U23B20155) and the Science and Technology Innovation Program of Hunan Province (grant number 2023RC1021). WW received support from the National Natural Science Foundation of China (grant numbers 42472027, 42342042) and the Natural Science Foundation of Hunan Province (grant number 2023JJ20063). We used Google’s Gemini to refine the language of the manuscript.

## Notes

### Competing Interest Statement

The authors have declared no competing interest.

