## Supplementary material for "Deep learning for mass extinction detection on fossilized phylogenies: power, limitations, and lessons for simulation-based phylodynamic inference": SI

### Supplementary Information

#### S1 Data Simulation Details

Given that our primary objective was to assess the feasibility of employing deep learning models for the prediction of mass extinction events, we introduced several constraints to manage the project's scope and computational demands. These measures were implemented to curtail the dimensionality of the parameter space, thereby mitigating the computational burden associated with both data simulation and model training. For example, we constrained the maximum fossilization rate to $0.05$, which served to prevent the generation of excessively large sampled phylogenies. Similarly, to reduce the computational time required for simulation and training, we constrained the number of tips (including both extant species and fossils, but excluding species that went extinct without being sampled as a fossil) in each phylogeny to a range of $200$ to $2,000$. We also restricted the origin time to a window between $80$ and $100$ time units, and set the sampling probability of extant species to $1$. It is important to note that while these constraints were pragmatic choices for this feasibility study, they are not inherent limitations of the overall approach. In theory, future work could address scenarios with different fossilization rates, origin times, and sampling probabilities by augmenting the volume of training data and expanding the model's parameter capacity. The complete simulation parameters are detailed in Table S1.

Instead of treating mass extinctions as instantaneous phenomena (i.e., a Dirac delta function), we modeled them as intervals of intense crisis by elevating the extinction rate within a randomly selected 1-Myr time bin. During each designated interval, the extinction rate was intensified by adding a value drawn from a uniform distribution between 0.4 and 1.0 to the existing background rate (Table S1). This range was chosen to model events of catastrophic severity, as the estimated magnitudes of several major mass extinctions in geological history fall within this interval (Bambach, 2006). An extinction rate of 0.4 within a 1-Myr time bin is equivalent to a 40% loss of species during that interval. This, in turn, implies that with a phylogeny of sufficient size and an accurately specified model, it is, in principle, possible to detect these events with accuracy. To create a clearer distinction between the simulated mass extinctions and background fluctuations in diversification, we meticulously calibrated our simulation parameters. We sought to establish a strong contrast between the two phenomena by deliberately maintaining low background rates, setting the maximum for both speciation and background extinction to 0.2 in an attempt to generate a more pronounced mass extinction signal.

The timing of these potential mass extinction intervals was restricted to occur between 20% and 80% of the tree's origin time (Table S1). This constraint was imposed because extinctions occurring too close to the beginning or end of the phylogeny are often difficult to detect (May et al., 2016). With an average total tree duration of 90 Myr, the eligible time window for a mass extinction was restricted to the central 60% of the tree's history (i.e., between the 20% and 80% marks), yielding an average of 54 eligible 1-Myr time bins. To ensure that approximately half of the simulated trees experienced mass extinctions, the probability for an event to occur in any single eligible time bin was set to approximately 0.009 (a 0.5 total probability distributed across 54 bins, i.e., $0.5/54 \approx 0.009$). This probabilistic assignment meant that any given tree could ultimately experience zero, one, or multiple mass extinction events. To develop and evaluate our model, we generated three distinct datasets using the simulation strategy outlined previously. A comprehensive training dataset was created, comprising 100,000 phylogenies. For performance assessment, we generated a validation dataset and an identical test dataset, each comprising 100,000 phylogenies.

To develop and evaluate our model, we generated three distinct datasets using the simulation strategy outlined previously. A comprehensive training dataset was created, comprising 100,000 phylogenies. For performance assessment, we generated a validation dataset and an identical test dataset, each comprising 100,000 phylogenies. The efficiency of the simulation process is a critical determinant in the application of simulation-based inference, particularly as substantial computational resources are frequently expended during the data generation phase. While a diverse array of simulation software exists (see Landis and Thompson, 2025), for this project, we utilized ReMASTER (Vaughan, 2024) in BEAST2 (Bouckaert et al., 2019) for the generation of all simulated datasets, as its high computational speed was essential for generating the vast number of phylogenies required for this study.

### S2 GNN Architecture Details

#### S2.1 Preprocessing Layers

The role of the preprocessing layer within a GNN architecture is to encode the features for each individual node (Figure S2A). This component is particularly effective when node attributes are represented by complex data types, such as images or text. In the present study, we implemented a relatively simple preprocessing layer. Our preliminary investigations, conducted on a simplified task, compared the impact of including versus omitting this layer (Section S3 and Figure S3). These evaluations revealed that its absence had only a marginal effect on performance (Figure S3), a finding likely attributable to the carefully selected and low-dimensional nature of our node features. Nevertheless, considering that our computational tasks were not excessively demanding in terms of resources, we elected to retain the preprocessing layer across all analytical undertakings to maintain a complete and robust architecture.

##### S2.1.1 GNN Layers

A GNN layer typically encompasses two fundamental operations: message computation and aggregation (Figure 1B and Figure S2B). First, message computation involves the transformation and feature extraction for each individual node. Subsequently, the aggregation step enables a given node to integrate the messages from itself and its neighbors, thereby forming its new feature representation. The specific implementations of these mechanisms differentiate various GNN architectures. In this study, we employed the Graph Attention Network (GAT) (Veličković et al., 2018). A significant advantage of the attention mechanism is that it obviates the need for manual specification of the aggregation strategy (e.g., summation or averaging); instead, the model autonomously learns to optimally weigh and integrate features from neighboring nodes (Figure 1B and Figure S2B). Furthermore, attention mechanisms can incorporate multiple “heads”, which can be conceptualized as distinct channels learning different feature aspects (Figure S2B). For instance, one head might focus on learning signals related to speciation rates, while another might concentrate on extinction rates. We utilized six such heads in our GAT layers.

To build a deep network, GNN architectures commonly incorporate multiple such layers. We implemented three GAT layers. To facilitate gradient flow and enhance learning in deeper architectures, we also introduced skip connections between different layers, a concept analogous to that employed in residual neural networks (He et al., 2016). The specific configuration of these connections is illustrated in (Figure 1A).

##### S2.1.2 Post-Process Layers

Finally, we employed a straightforward post-processing layer, which was designed to transform the LSTM network's output into a format suitable for our specific classification or regression tasks.

### S3 Ablation Studies

To assess how different architectural components contribute to the model performance, we conducted a series of ablation studies using our simulated datasets. The performance of the complete, fully-integrated model is detailed in Section 3.7. We then systematically evaluated the impact of removing individual components—specifically, the preprocessing layers, the Virtual Nodes, the time bin pooling mechanism, and the LSTM layer (Table S2). The performance of each of these modified architectures is presented in Figure S3, illustrating the relative importance of each component for accurate extinction detection.

### S4 Bayesian Benchmark

For our Bayesian benchmark, we analyzed phylogenies using a SFBD model with horseshoe Markov random field prior implemented in `RevBayes`. The model's parameters and prior distributions were configured to be identical to those used in the data simulation process, ensuring a consistent comparison.

We discretized time into intervals of 1 Myr, starting from the tree's origin. These intervals served as potential points for both background rate shifts and the mass extinction event. We set a prior expectation of 0.5 mass extinction events for any given tree. The window for a potential mass extinction was restricted to the period between 20 and 70 Ma. Within this window, the prior probability of a mass extinction occurring at any specific 1-Myr interval was defined is $0.5/50 = 0.01$, with the species survival probability during the event drawn from a uniform distribution between 0.0 and 0.5. The prior probability of a mass extinction occurring outside the 20–70 Ma window was zero.

Each Markov chain Monte Carlo (MCMC) analysis was run for 200,000 iterations, with the initial 40,000 iterations discarded as burn-in. We assessed convergence using ESS. For the 200 trees that were incorrectly classified by the deep learning model, only a single tree had an analysis with an ESS value below 200. For the 200 randomly selected trees, only five analyses yielded an ESS below 200. Given the small size of our benchmark dataset, we decided to retain all 200 analyses in our results without excluding those with lower ESS values.

Within a Bayesian framework, the estimated time of mass extinction is identified as the time point associated with the highest Bayes factor. Subsequently, the inferred extinction proportion is calculated by combining the background extinction rate observed at this peak Bayes factor time point with the complement of the extinction probability (i.e., one minus the survival probability). To determine this survival probability, we compute the average of all survival probabilities less than one from time points located near the maximum Bayes factor, specifically including those where the $2 \log \mathrm{BF0}$ is greater than zero.

### S5 Detailed Model Tuning and Performance Exploration

After refining the dataset, which elevated the model's accuracy to ˜70%, we conducted a series of systematic experiments to determine if further improvements were being constrained by the model's configuration, architecture, or the size of the training data (Table S2 and Figure S4).

#### S5.1 Model Complexity

First, we addressed model complexity as a potential limiting factor. To test this, we doubled the model's key parameters, creating a substantially more complex architecture (model 3 in Table S2). This larger model, however, performed similarly to the original, achieving nearly identical classification accuracy (Figure S4). This result suggested that performance was not being constrained by the model's complexity, and that simply increasing its size further would be unlikely to yield better results.

#### S5.2 Hyperparameter Tuning

Next, we explored the possibility of suboptimal hyperparameter settings. We conducted a series of experiments adjusting the model's dropout probabilities and learning rates (Goodfellow et al., 2016) (models 4-8 in Table S2). However, these modifications did not yield any significant improvement in accuracy (Figure S4). The only notable outcome was the confirmation that an excessively high learning rate (e.g., 0.01) severely impaired the model's ability to learn, reinforcing that our initial settings were appropriate (Figure S4).

#### S5.3 Alternative Network Architecture

We also explored an alternative neural network architecture. We replaced the model's core LSTM component with a simplified Transformer block (model 9 in Table S2), a design known for its power in sequence analysis (Vaswani et al., 2017). This fundamental architectural change, however, not only failed to improve performance but, in fact, led to a degradation in predictive accuracy (Figure S4).

#### S5.4 Training Data Size

Finally, we investigated whether the model was data-limited. We tripled the size of our training dataset, expanding it from 100,000 to 300,000 simulated phylogenies (model 10 in Table S2). Despite this substantial increase in data volume, the model's accuracy remained static (Figure S4). Having systematically ruled out hyperparameters, model complexity, architecture, and data volume as the source of the performance plateau, we concluded that a more fundamental aspect of the data or model structure was at play.

#### S6 Detailed Analysis of Factors Influencing Mass Extinction Detection Accuracy

To gain a deeper understanding of the model's performance, we evaluated the relationship between its accuracy and scores against a range of phylogenetic parameters. The observed relationships were consistent across our training, validation, and test sets.

The model's ability to correctly identify the presence or absence of a mass extinction improved with a greater number of tips (Figure S18) and fossils (Figure S19). In both cases, performance increased steadily before plateauing around 1,000 tips (Figure S18) or 200 fossils (Figure S19), respectively. This suggests a point of diminishing returns where additional data provides little new information for the model. Notably, the performance boost from more tips or fossils was more pronounced when detecting a true mass extinction compared to confirming its absence.

A more perplexing pattern emerged in relation to the number of extant species. For both phylogenies with and without mass extinctions, the model's accuracy followed a U-shaped curve, initially decreasing as extant species increased, reaching a minimum at around 200 extant species, and then improving again (Figure S20). We observed that when the number of extant species was very low, the model was prone to misclassifying trees that had not experienced a mass extinction as having had one (Figure S20). We attribute this counterintuitive trend to a sampling bias in our simulation protocol. A large fraction of our simulated phylogenies feature continuously increasing diversity, leading to a high number of extant species. Consequently, phylogenies with low extant diversity are disproportionately those that have suffered a mass extinction. The model appears to have learned this spurious correlation, associating low extant richness with mass extinction, even though no direct causal link exists. This artifact likely explains the misclassification of the phylogeny in Figure S10 and suggests a need to augment our training data with more phylogenies that feature low species richness without a history of mass extinction.

The proportion of species eliminated during a mass extinction had a nuanced effect on the model's performance. Initially, as the extinction proportion grew, the model's accuracy increased, reaching its peak when approximately 67% of the species were wiped out (Figure S21). However, accuracy then declined at even higher extinction levels. This drop-off appears to be an artifact of our data generation and filtering process. Although we simulated extinction magnitudes from a uniform distribution between 50% and 100%, the compounding effect of background extinction should have produced scenarios with even greater total species loss. It is likely that our filtering criterion, which required phylogenies to retain more than 200 tips, systematically removed these most extreme extinction events from our final dataset, thus creating an underrepresentation of such cases.

In contrast to the complex influence of extinction proportion, the effect of the number of extinct species was more direct. Both the model's extinction score and its accuracy improved as more species were lost. This trend continued until the number of extinctions surpassed 100, at which point the performance plateaued (Figure S22). This suggests that the loss of 100 or more species provides a signal that is strong and unambiguous enough for the model to reliably detect.

Finally, the pre-extinction diversity and timing of the event were critical. Performance improved with higher pre-extinction diversity, stabilizing once this number exceeded 200 lineages (Figure S23). The timing of the mass extinction was also highly influential: extinctions occurring later in a tree's history were detected with much higher accuracy (Figure S24). This trend continued to the edge of our simulated window (80% of total time). Of particular note, when a mass extinction occurred early in a tree's history (around 30% of its total duration), the model's score and accuracy hovered near 0.5. This indicates that the model has virtually no predictive power for these events, as their phylogenetic signal becomes significantly attenuated over long evolutionary timescales.

### S7 Detailed Robustness Assessment

To quantitatively assess the model's robustness to phylogenetic uncertainty, we subjected a set of 2,000 phylogenies (1,000 with a mass extinction and 1,000 without) to a series of perturbations designed to mimic the uncertainty inherent in Bayesian phylogenetic inference.

#### S7.1 Internal Node Time Perturbations

To assess the model's sensitivity to uncertainty in the timing of evolutionary events, we systematically altered the ages of the internal nodes in our test phylogenies. Our approach was designed to closely mimic the process of Bayesian phylogenetic inference by emulating the proposal mechanisms used in the software `BEAST2`. Specifically, we employed the `BactrianNodeOperator`, which randomly selects an internal node and proposes a new age for it while keeping the tree topology. We configured this operator to use a bactrian distribution, a setting that encourages the proposed new node ages to be substantially different from their previous values, thereby ensuring a rigorous test of the model. Furthermore, to understand the impact of the proposal size, we systematically adjusted the `scaleFactor` parameter of the `BactrianNodeOperator`, which controls the size of the age changes, evaluating the model's response at settings of 0.1, 0.3, 1.0, and 2.0 (Figure S27).

We assigned a placeholder molecular character to each species, which is a necessary step to run the MCMC machinery. The analysis was configured to use only the `BactrianNodeOperator`, ensuring that only node ages were modified. Furthermore, we used a simple Yule tree prior with a uniform distribution for the birth rate, a technical setup that guarantees every proposed change to a node's age is accepted. The MCMC for each phylogeny began with the true, un-altered tree. To ensure that trees of different sizes were subjected to a comparable degree of disruption, we scaled the number of MCMC iterations to be equal to the number of tips in the tree. From this process, we saved 10 perturbed trees at regular intervals (specifically, after every `floor(number of tips / 10)` iterations), creating a gradient of disruption that ranged from approximately 10% to 100% of the internal nodes being modified. This proportion was subsequently recorded as an 'index' of perturbation in Figure S27.

Our results show that the deep learning model is remarkably robust to this type of uncertainty. Although the model's mass extinction score for the perturbed trees gradually diverged from the original score as the perturbations became more severe (i.e., with larger `scaleFactor` values or more modified nodes), the deviation was not substantial (Figure S25). Because this is a classification task, the final prediction—whether a mass extinction occurred or not—remained highly stable. Even under the most extreme scenario we tested, the vast majority of perturbed phylogenies yielded the same prediction as the original, unperturbed tree (Figure S25). This high degree of stability indicates that the model is robust and has not overfitted to the specific node times in the training data.

This pronounced robustness is likely attributable to the fact that the absolute magnitude of the temporal perturbations was ultimately limited. Since our phylogenies contain a minimum of 200 species, a large proportion of the branches are inherently short, constraining how much any single node's age can be shifted without violating the tree's topology. Our test data confirmed this: even under the most significant disruptions, the average time shift per node remained small, typically falling between 0 and 0.4 million years, and the maximum observed shift remained minimal (Figure S27). Nevertheless, the model was not entirely insensitive to these modifications. As the magnitude of the temporal perturbations increased, the model's mass extinction score progressively diverged from its original value, demonstrating a predictable response to the introduced uncertainty (Figure S27).

#### S7.2 Topology Perturbations

Following the same approach used for node ages, we next tested the model's robustness to uncertainty in the tree's branching structure, or topology. We introduced perturbations using the `Exchange` operator from `BEAST2`, a method that performs a nearest-neighbor interchange (NNI) to alter the topological connections between lineages while preserving the original ages of the internal nodes. The MCMC setup was identical to that used for the node-time perturbations, with the only differences being the operator employed and the sampling frequency. For this experiment, we saved five perturbed trees at regular intervals—specifically, after every `floor(number of tips / 5)` iterations—to generate a gradient of increasing topological disruption. This proportion was subsequently recorded as an 'index' of perturbation in Figure S28.

The model demonstrated substantial robustness to these changes. As the number of topological modifications increased, the model's mass extinction score progressively diverged from its original value. However, the final classification remained highly stable. Even under the most extreme perturbation scenario, the predictions for the majority of the altered phylogenies were identical to those for the original, unperturbed trees (Figure S26).

To quantify the extent of the topological changes more formally, we calculated the Robinson-Foulds (RF) distance between each perturbed tree and its original simulated counterpart. We observed a clear trend: as the RF distance increased, the model's score diverged further from its initial value (Figure S28). However, even when the RF distance approached 0.5—indicating a significant topological rearrangement—the model often still correctly predicted the presence of a mass extinction (Figure S28).

This relative insensitivity to topology suggests that the model's classification relies on features that are not strictly dependent on the fine details of the branching pattern. This can be explained by the nature of the underlying skyline FBD model used to simulate the data. In this framework, the key signals of a shift in diversification dynamics are the aggregate counts of originations and extinctions within discrete time intervals. Consequently, as long as the total number of these events per interval remains consistent, the model's inference is largely unaffected by where on the phylogenetic tree these originations and extinctions occur. However, this reasoning is predicated on the assumption of a homogeneous diversification rate across all lineages. If this assumption were relaxed—for instance, by using a multi-type model where different lineages have distinct diversification rates—the tree's topology would likely become a much more significant factor in the model's performance. Furthermore, because the simulated mass extinctions were non-selective (i.e., random), the precise topological relationships among lineages are inherently less informative for detecting the event.

##### S7.3 Combined Internal Node Time and Topological Perturbations

To further explore the impact of perturbations on both node ages and tree topology, we employed a combined strategy utilizing the `BactrianNodeOperator` and the `Exchange` operator. This combination allowed for simultaneous modification of internal node ages and the tree's topology. Both operators were assigned an equal weight of 1. For the `BactrianNodeOperator`, the `scaleFactor` was specifically set to 1.0. The overall MCMC configuration largely mirrored that of the previously described experiment, with the primary distinction being the concurrent application of these two operators. In this setup, the MCMC chain was run for a total number of iterations equal to twice the number of tips in the tree. The tree state at the conclusion of this MCMC run was then designated as the final perturbed tree.

Consistent with our previous findings, the model's predictions remained remarkably stable overall, with the results from the perturbed trees not deviating substantially from the original analyses (Figure 4).

#### S8 Detailed Model Mismatch Analysis

We evaluated the model's performance on two new datasets simulated under conditions that were not represented in the training data.

##### S8.1 Mismatch with Increasing Phylogenetic Tree Size

We simulated a new dataset of larger trees, constraining the number of tips to be between 2,200 and 4,000, while all other simulation parameters remained identical to those used for the original training data. This process generated two sets of 3,000 trees: one intended to have no mass extinction and another intended to include one. An examination of the resulting phylogenies confirmed that all 3,000 trees simulated without a mass extinction event were indeed free of them. For the set intended to include a mass extinction, 1,975 of the 3,000 simulations successfully produced a phylogeny with a single, distinct mass extinction event. Notably, this success rate ($1975/3000 \approx 66\%$) is higher than that observed in the generation of the original training data (55%). This is likely because larger trees, on average, have higher standing diversity. A greater number of species at the time of the event provides more lineages subject to extinction, making it more probable that the stochastic simulation process will result in a realized extinction event that meets our dual criteria of both high proportional loss and a significant absolute number of extinctions.

When we applied our trained model to this new set of larger trees, we observed a mixed but informative pattern of performance. The model's accuracy in detecting the presence of a mass extinction improved (Figure S5K). This indicates that the model can be generalized to larger trees to some extent, a robustness we attribute to two key aspects of our design: the time-bin pooling strategy and the inherent properties of the GNN architecture. First, because the number of time bins is fixed, increasing the number of tips in a tree simply increases the number of lineages within each bin. Our pooling strategy, however, averages the information from all lineages within a bin. Consequently, the informational input from a bin with ten lineages is not dramatically different from that of a bin with one, which dampens the effect of tree size. Second, in the GNN, each node's features are updated based only on its immediate neighbors. In a tree structure, this neighborhood is fixed (an internal node always has three neighbors, and a tip has one), so increasing the overall size of the tree does not alter the local computational structure for any given node in GNN.

Nevertheless, this scalability has its limits. The performance of any deep learning model is fundamentally tied to its training data, and we anticipate that its accuracy would degrade on exceptionally large trees with tens of thousands of tips or more. We did not extend our tests to such scales, primarily because generating such large phylogenies is difficult under our current simulation parameters without producing unrealistic outlier trees. For future work, a promising approach to generate larger, more stable trees would be to increase the fossilization rate while maintaining the same birth and death rates.

#### S8.2 Mismatch with Increasing Diversification Rate

We next evaluated the model's performance under conditions of elevated diversification rates. To do this, we simulated a new dataset by increasing the range for both birth and death rates to

0.25–0.45 per time bin. As before, we generated 3,000 trees intended to be free of mass extinctions and 3,000 trees intended to contain one. The simulation outcomes highlighted a critical issue: of the 3,000 control trees, 585 were flagged as having a mass extinction, while 2,584 of the 3,000 target trees contained at least one mass extinction event. Furthermore, among the trees with extinctions, 903 contained multiple mass extinction events. This high frequency suggests that our definition of a mass extinction becomes less robust when background extinction rates are high, as stochastic fluctuations in the background extinction rate can be easily misidentified as a true mass extinction event under these conditions.

When we applied our model to this high-rate dataset, its performance was mixed. The model's accuracy in detecting trees that genuinely contained a mass extinction was higher (Figure S5L). However, it struggled to correctly classify trees that did not have a mass extinction, showing a high false-positive rate (Figure S5L). This is likely because the presence of multiple, strong extinction signals in many trees makes them easier to detect, while the elevated background extinction rate simultaneously blurs the distinction between normal and catastrophic mass extinction levels. This result clearly indicates that the model's predictive capabilities are substantially reduced when confronted with diversification dynamics outside the range of its training data.

### S9 Data Binning and Trend Fitting

To reveal underlying trends within the large and sparse dataset, we employed a binning and averaging method. The dataset, comprising several hundred thousand points, was first sorted by the independent variable (x-axis) and then partitioned into 1,000 sequential, equally-sized bins. The mean of the dependent y-values was then calculated for each bin to represent its central tendency.

Table S1: Parameters for the initial simulation and refinement simulation.

| **Parameter** | **Value** |
|---|---|
| **Horseshoe Markov Random Field Skyline FBD** | |
| Origin Time ($t_{\text{ori}}$) | $U(80, 100)$ |
| Number of Time Bins | $\lceil t_{\text{ori}} \rceil$ |
| Initial Birth Rates ($\lambda_0$) | $\min(0.005, 0.2 \cdot \text{Beta}(1.5, 1.5))$ |
| Initial Death Rates ($\mu_0$) | $\min(0.005, 0.2 \cdot \text{Beta}(1.5, 1.5))$ |
| Initial Fossilization Rates ($\psi_0$) | $\min(0.005, 0.05 \cdot \text{Beta}(1.5, 1.5))$ |
| Global Scale hyperparameter of Birth Rates ($\zeta_\lambda$) | $0.0021$ |
| Global Scale hyperparameter of Death Rates ($\zeta_\mu$) | $0.0021$ |
| Global Scale hyperparameter of Fossilization Rates ($\zeta_\psi$) | $0.0021$ |
| Global Scale of Birth Rates ($\gamma_\lambda$) | $\text{HalfCauchy}(0, \zeta_\lambda)$ |
| Global Scale of Death Rates ($\gamma_\mu$) | $\text{HalfCauchy}(0, \zeta_\mu)$ |
| Global Scale of Fossilization Rates ($\gamma_\psi$) | $\text{HalfCauchy}(0, \zeta_\psi)$ |
| Local Scale of Birth Rates $\sigma_\lambda$ | $\text{HalfCauchy}(0, 1)$ |
| Local Scale of Death Rates $\sigma_\mu$ | $\text{HalfCauchy}(0, 1)$ |
| Local Scale of Fossilization Rates $\sigma_\psi$ | $\text{HalfCauchy}(0, 1)$ |
| Extant Sampling Probability ($\rho$) | $1$ |
| **Initial Simulation Setting** | |
| Time of Mass Extinction | $U(0.2 \cdot t_{\text{ori}}, 0.8 \cdot t_{\text{ori}})$ |
| The Probability of A Mass Extinction Occurring in One Time Bin | $0.006$ |
| Extinction Rate of Mass Extinction $\mu_{\text{ME}}$ | $U(0.5, 1.0)$ |
| **Refinement Simulation Setting** | |
| Time of Mass Extinction | $\text{RandomInteger}(\text{round}(0.3 \cdot t_{\text{ori}}), \text{round}(0.8 \cdot t_{\text{ori}}))$ |
| Number of Mass Extinction | $0/1$ |
| Extinction Rate of Mass Extinction $\mu_{\text{ME}}$ | $U(0.4, 1.0)$ |

Table S2: Summary of the parameters and training data used for each model configuration. All models uniformly utilized a weight decay of 0.0001, a batch size of 128, and an output dimension of 1.

| Section | Model index | Training Data | learning rate | dropout probability | preprocess dim | GAT layers | GAT hidden dim | GAT heads | LSTM layers | LSTM hidden dim | postprocess dim |
|---|---|---|---|---|---|---|---|---|---|---|---|
| 3.1 | 1 | 100000 (Initial simulation setting) | 0.001 | 0.5 | 256 | 3 | 64 | 3 | 1 | 64 | 64 |
| 3.2 | 2 | 100000 (refinement simulation setting) | 0.001 | 0.5 | 256 | 3 | 64 | 3 | 1 | 64 | 64 |
| 3.3 | 3 | 100000 (refinement simulation setting) | 0.001 | 0.5 | 256 | 3 | 64 | 6 | 2 | 128 | 64 |
| 3.3 | 4 | 100000 (refinement simulation setting) | 0.001 | 0.3 | 256 | 3 | 64 | 6 | 2 | 128 | 64 |
| 3.3 | 5 | 100000 (refinement simulation setting) | 0.01 | 0.5 | 256 | 3 | 64 | 6 | 2 | 128 | 64 |
| 3.3 | 6 | 100000 (refinement simulation setting) | 0.03 | 0.3 | 256 | 3 | 64 | 6 | 2 | 128 | 64 |
| 3.3 | 7 | 100000 (refinement simulation setting) | 0.0001 | 0.5 | 256 | 3 | 64 | 6 | 2 | 128 | 64 |
| 3.3 | 8 | 100000 (refinement simulation setting) | 0.0001 | 0.3 | 256 | 3 | 64 | 6 | 2 | 128 | 64 |
| 3.3 | 9 | 100000 (refinement simulation setting) | 0.001 | 0.5 | 256 | 3 | 64 | 3 | 3 layers, 3 heads, 1024 dim | | 64 |
| 3.3 | 10 | 300000 (refinement simulation setting) | 0.001 | 0.5 | 256 | 3 | 64 | 6 | 2 | 128 | 64 |
| 3.7 | 11 | 300000 (refinement simulation setting, redefine mass extinction) | 0.001 | 0.3 | 256 | 3 | 64 | 6 | 2 | 256 | 64 |
| 2.3.3 | 12 | 300000 (refinement simulation setting, redefine mass extinction) | 0.001 | 0.3 | - | 3 | 64 | 6 | 2 | 256 | 64 |
| 2.3.5 | 13 | 300000 (refinement simulation setting, redefine mass extinction, no virtual nodes) | 0.001 | 0.3 | 256 | 3 | 64 | 6 | 2 | 256 | 64 |
| 2.3.6 | 14 | 300000 (refinement simulation setting, redefine mass extinction, global mean pooling) | 0.001 | 0.3 | 256 | 3 | 64 | 6 | - | - | 64 |
| 2.3.7 | 15 | 300000 (refinement simulation setting, redefine mass extinction) | 0.001 | 0.3 | 256 | 3 | 64 | 6 | 2 MLP (first: 38885-1024, second: 1024-512) | | 64 |

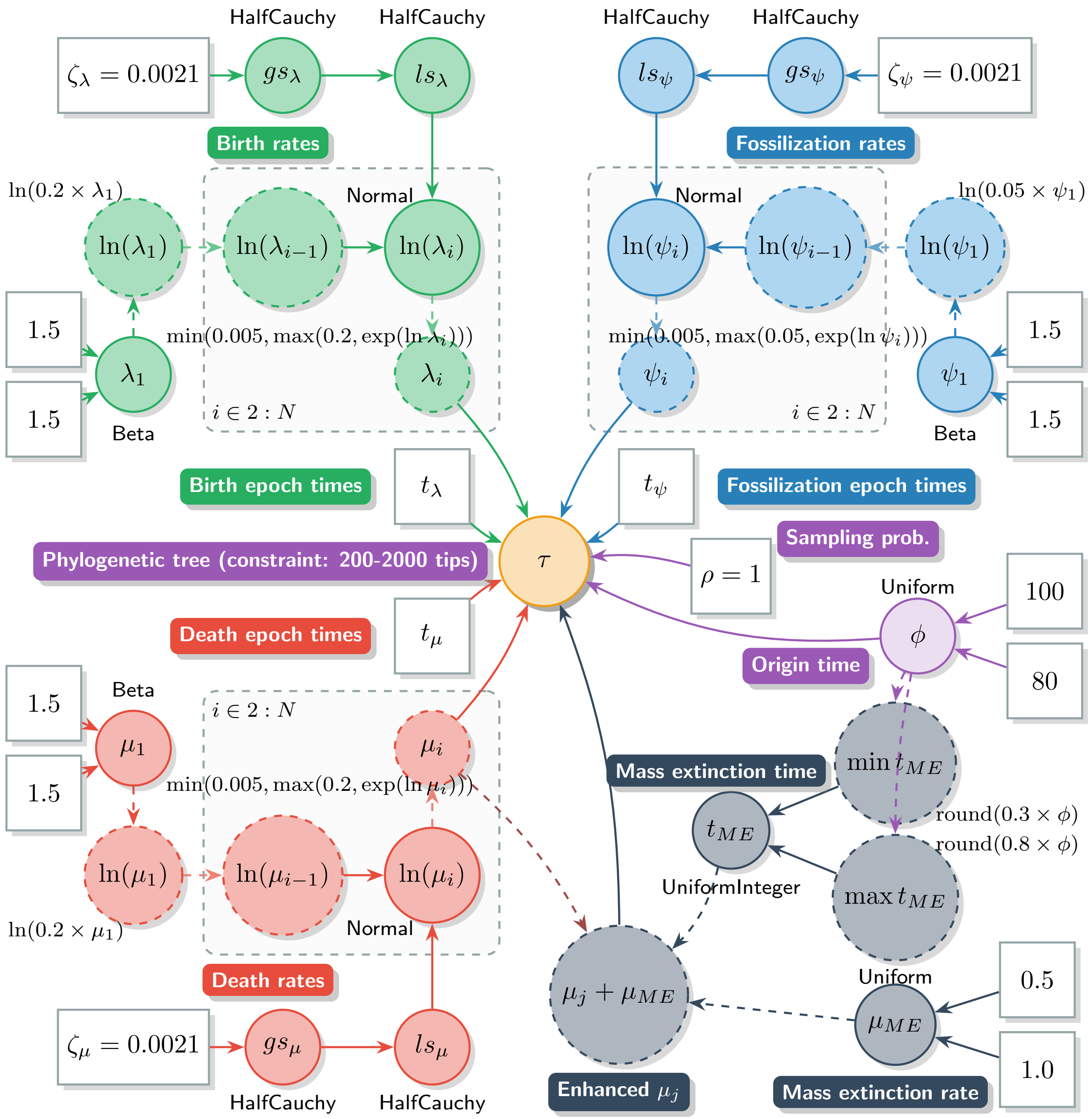


Figure S1: A diagram illustrating the simulation workflow for the skyline FBD model. This model incorporates a horseshoe Markov random field prior and is constrained to simulate the occurrence of a single mass extinction event (refinement simulation setting).

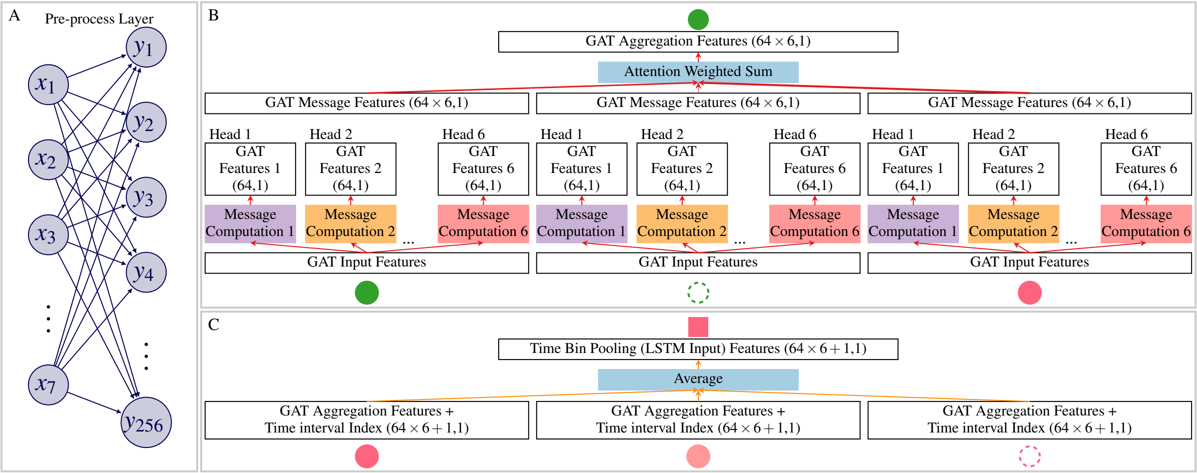


Figure S2: Key components of the model architecture. (A) A schematic of the pre-processing layer. (B) A schematic of a GAT layer, detailing the message computation and aggregation steps. (C) A schematic of the time bin pooling mechanism.

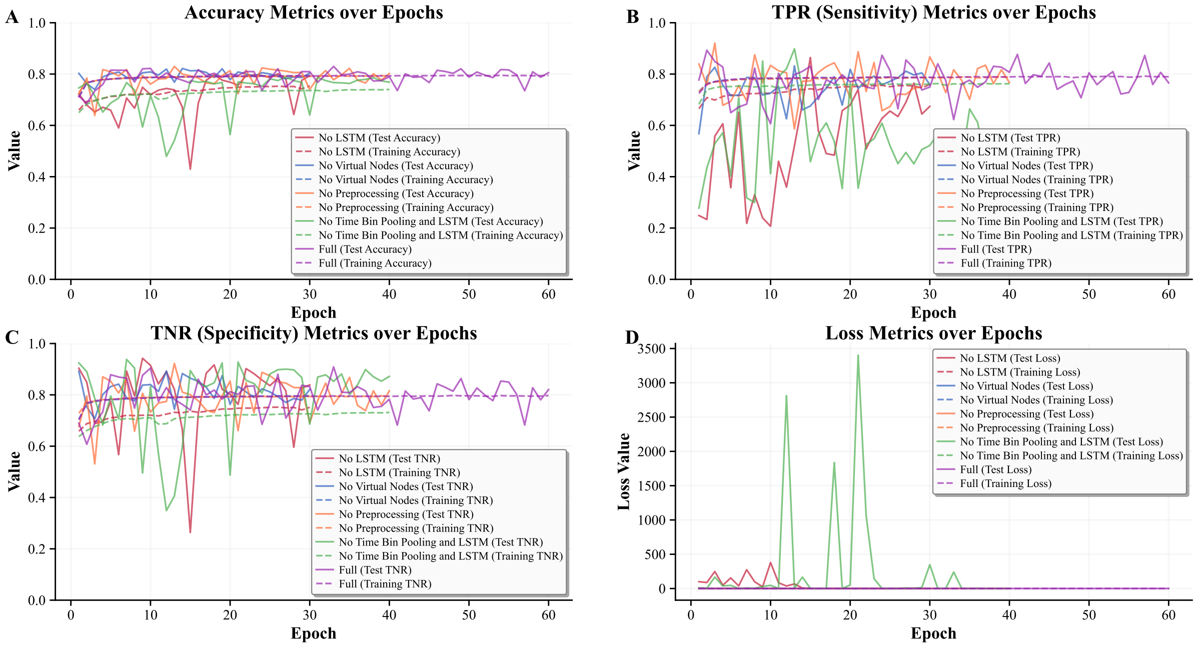


Figure S3: An ablation study assessing the contribution of individual model components to the detection of mass extinctions. This figure compares key performance metrics—including accuracy, TPR, TNR, and loss—between the full model and several variants, each with a specific component removed: the LSTM, the virtual node, the preprocessing layers, or the time bin pooling and LSTM.

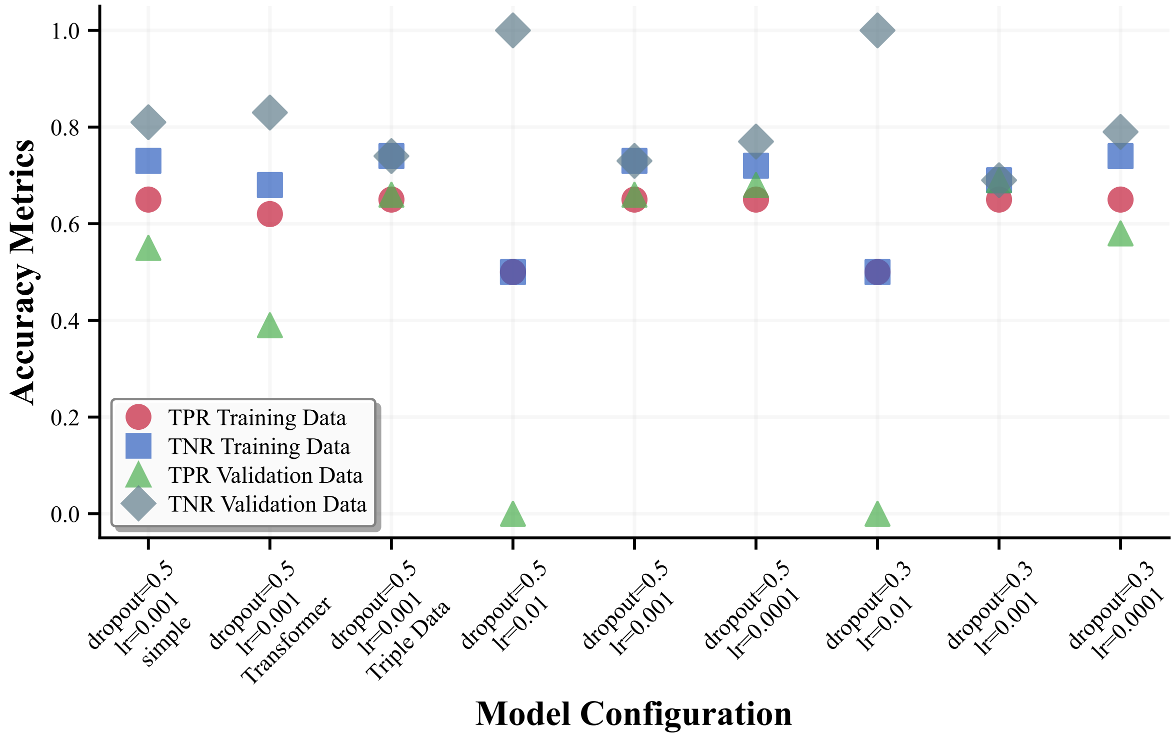


Figure S4: A sensitivity analysis of the model's performance evaluated on simulated phylogenies, both with and without mass extinction events. This figure compares the TPR and TNR across several experimental conditions, including: adjustments to the learning rate and dropout probability, a reduction in the number of model parameters, the replacement of the LSTM architecture with a Transformer, and a threefold increase in the size of the training dataset.

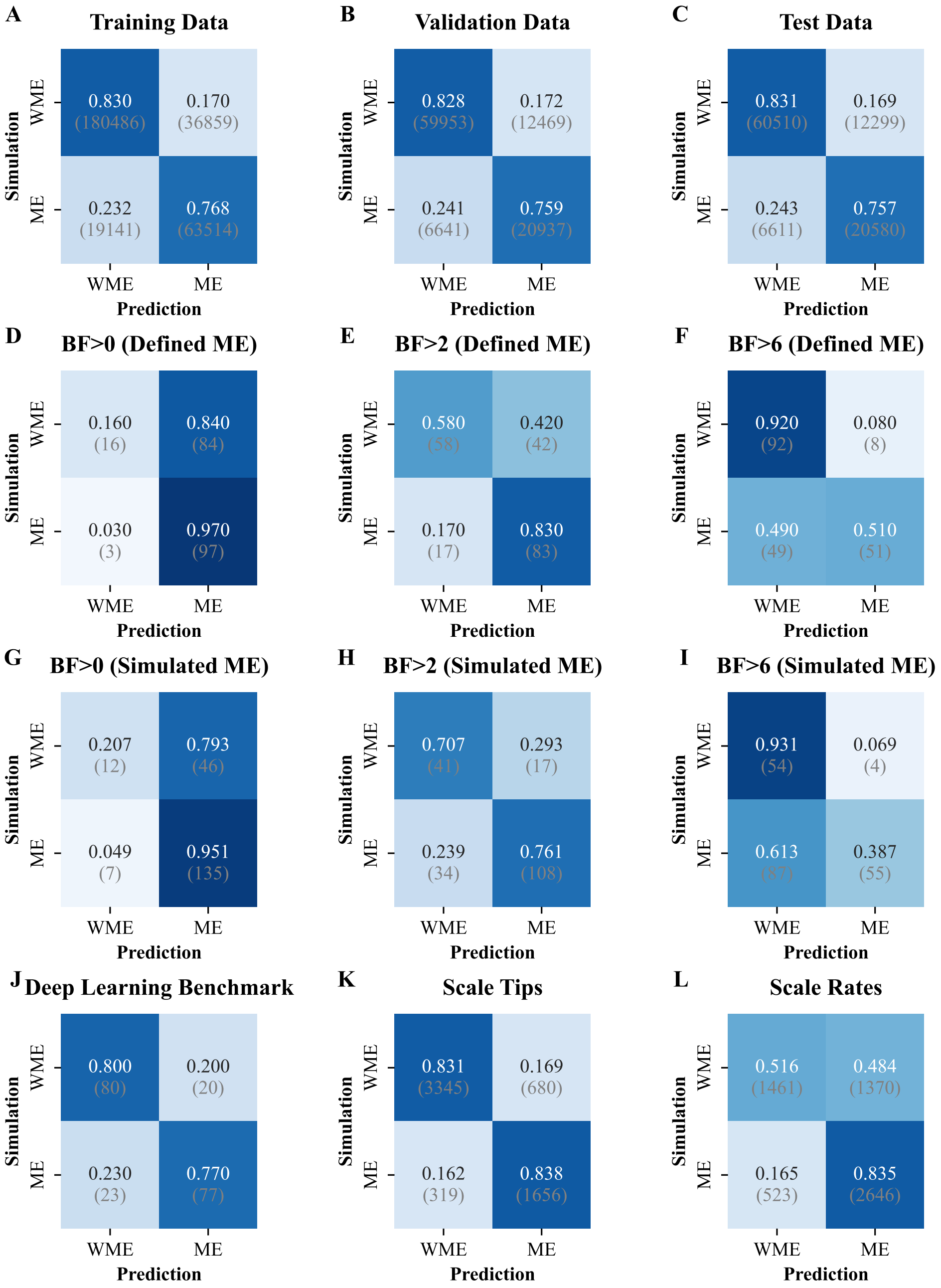


Figure S5: Confusion matrices illustrating the comparative classification accuracy of the deep learning and Bayesian methods across various datasets and conditions. The subplots represent the following analyses: (A-C) Performance of the final deep learning model on the redefined training, validation, and test datasets, respectively. (D-F) Performance of the Bayesian benchmark analysis using Bayes factor thresholds of $> 0$, $> 2$, and $> 6$, respectively, on phylogenies labeled according to our redefined criteria for a mass extinction. (G-I) Performance of the Bayesian benchmark with the same Bayes factor thresholds, but on phylogenies labeled according to the original simulation parameters (i.e., whether an event was intended to occur). (J) Performance of the deep learning model on the randomly selected benchmark dataset used for direct comparison with the Bayesian method. (K-L) Performance of the deep learning model under model mismatch scenarios, when tested on phylogenies with a larger number of tips (K) and higher background diversification rates (L) than seen in the training data. ME: Mass Extinction; WME: Without Mass Extinction. Numbers indicate the proportion of predictions for each true class, with the absolute counts of phylogenies in parentheses.

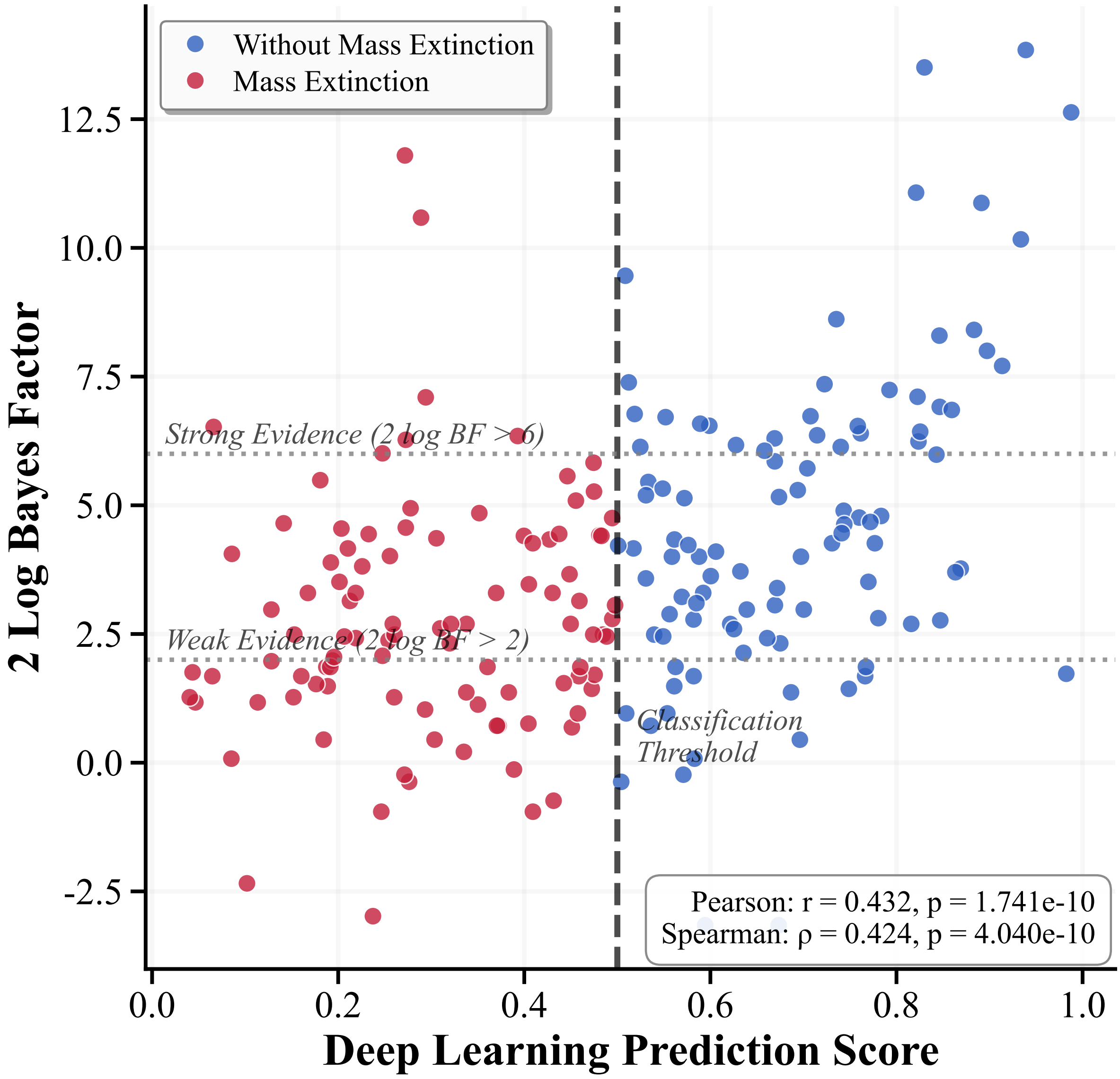


Figure S6: A scatter plot of the deep learning model's extinction prediction score versus the Bayes factor from the Bayesian method. The plot shows data from 100 phylogenies where a mass extinction occurred but was not detected by the model (false negatives), and 100 phylogenies where no extinction occurred but was incorrectly detected by the model (false positives).

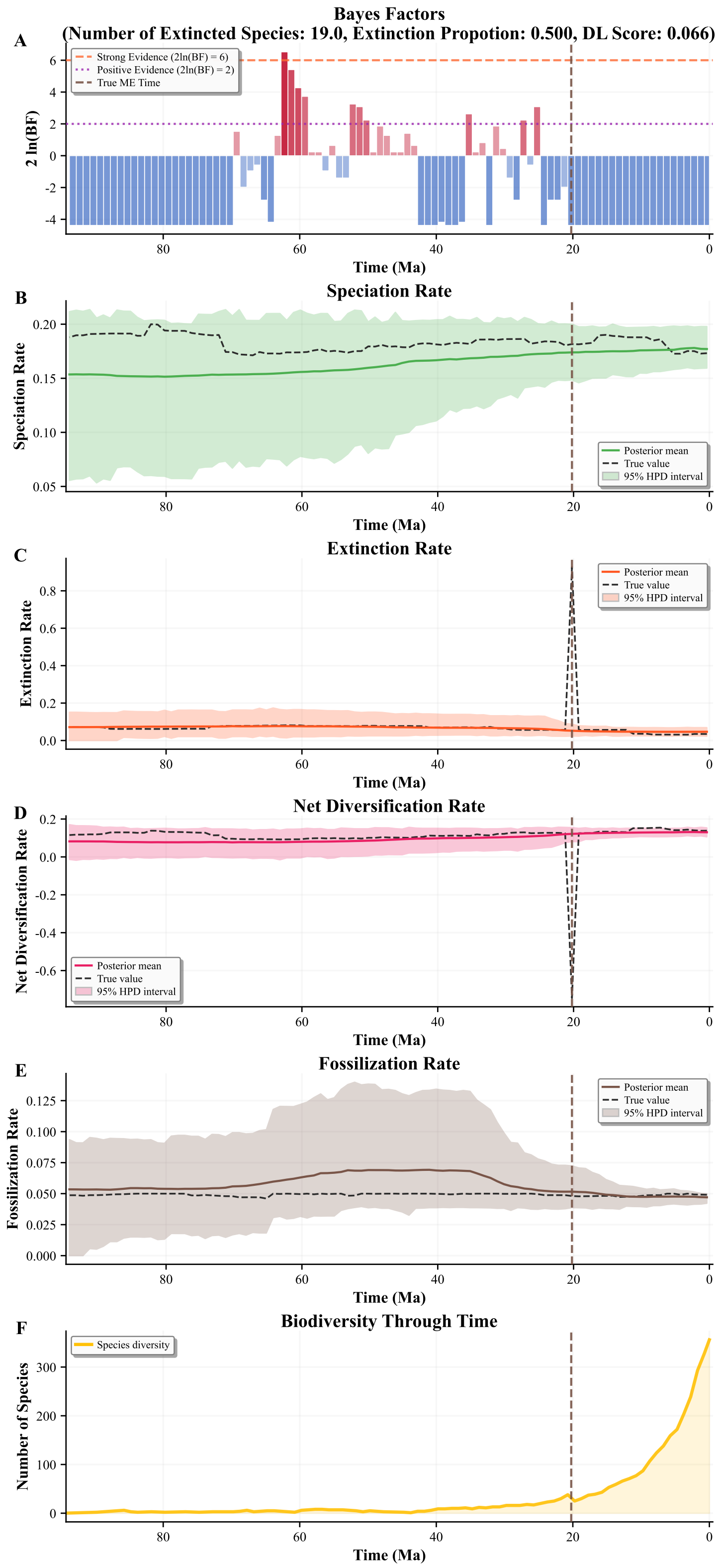


Figure S7: Deep learning and Bayesian inference for a simulated evolutionary history. (A) deep learning score and Bayes factor, (B-E) key evolutionary rates (speciation, extinction, diversification, and fossilization), and (F) the overall trajectory of species diversity through time.

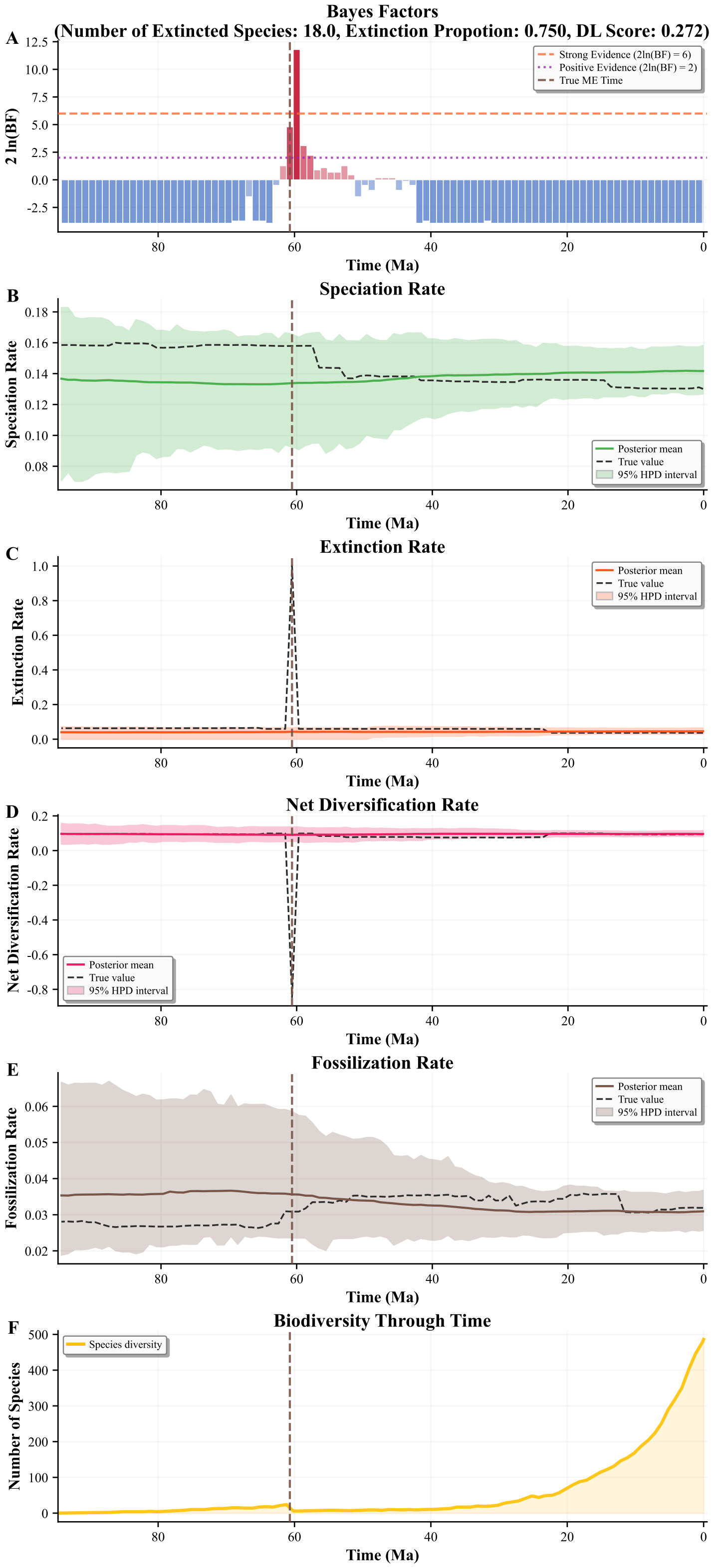


Figure S8: Deep learning and Bayesian inference for a simulated evolutionary history. (A) deep learning score and Bayes factor, (B-E) key evolutionary rates (speciation, extinction, diversification, and fossilization), and (F) the overall trajectory of species diversity through time.

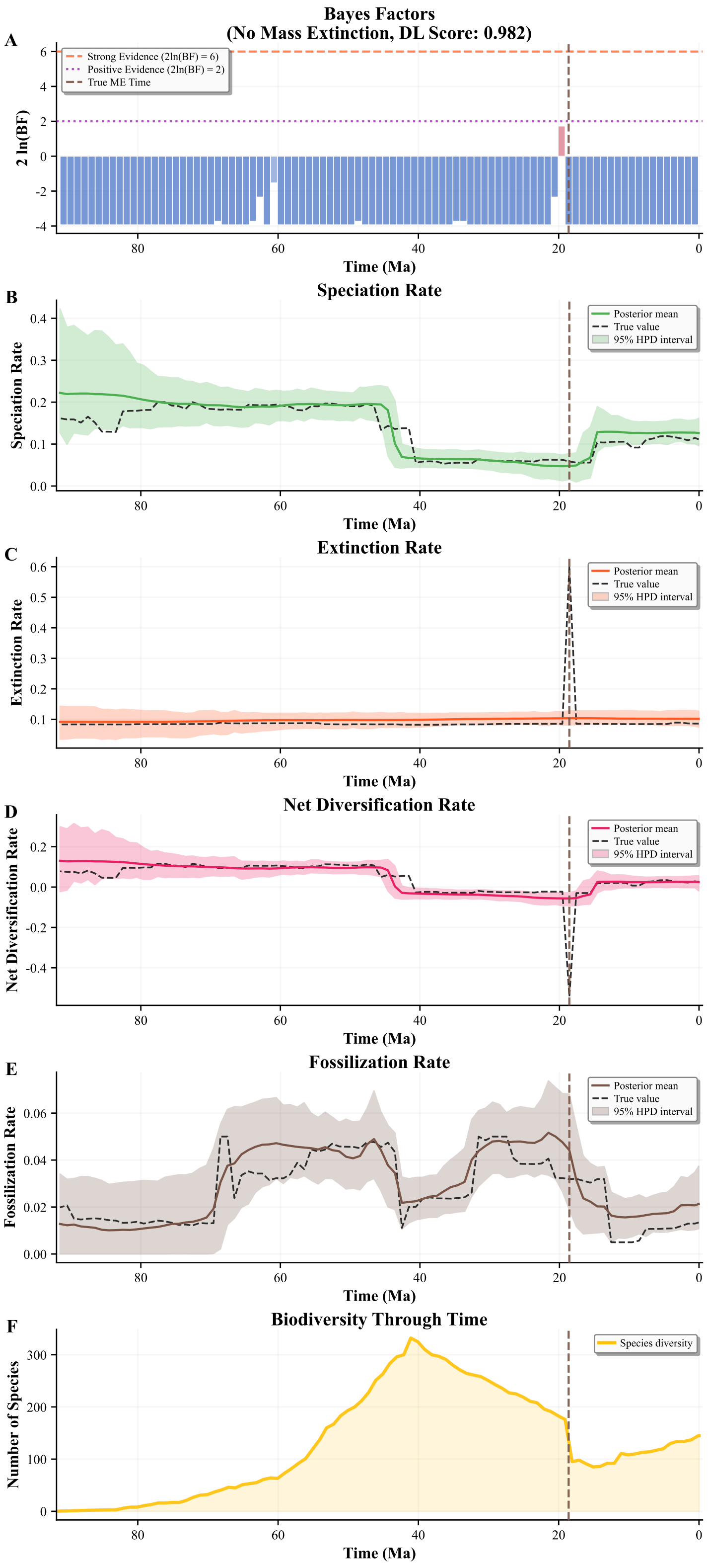


Figure S9: Deep learning and Bayesian inference for a simulated evolutionary history. (A) deep learning score and Bayes factor, (B-E) key evolutionary rates (speciation, extinction, diversification, and fossilization), and (F) the overall trajectory of species diversity through time.

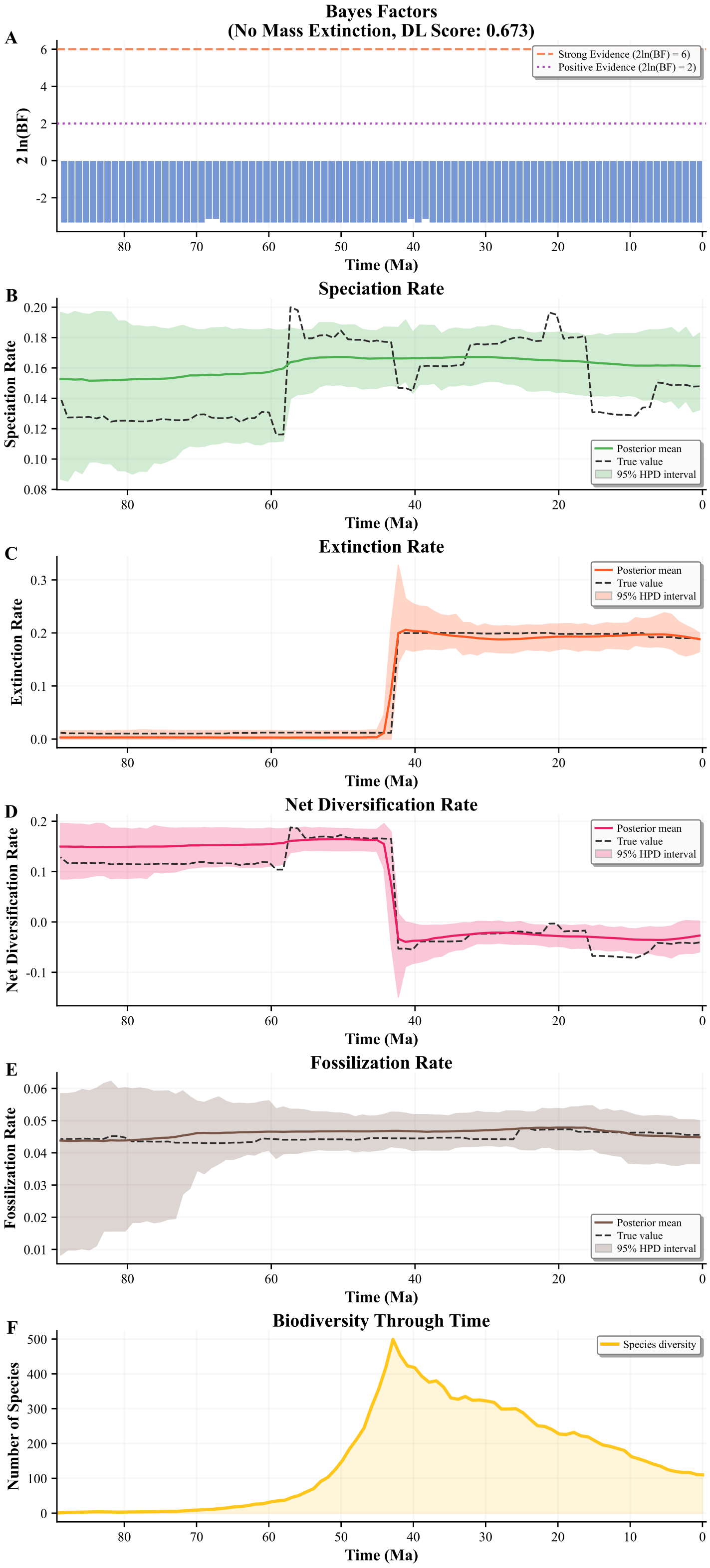


Figure S10: Deep learning and Bayesian inference for a simulated evolutionary history. (A) deep learning score and Bayes factor, (B-E) key evolutionary rates (speciation, extinction, diversification, and fossilization), and (F) the overall trajectory of species diversity through time.

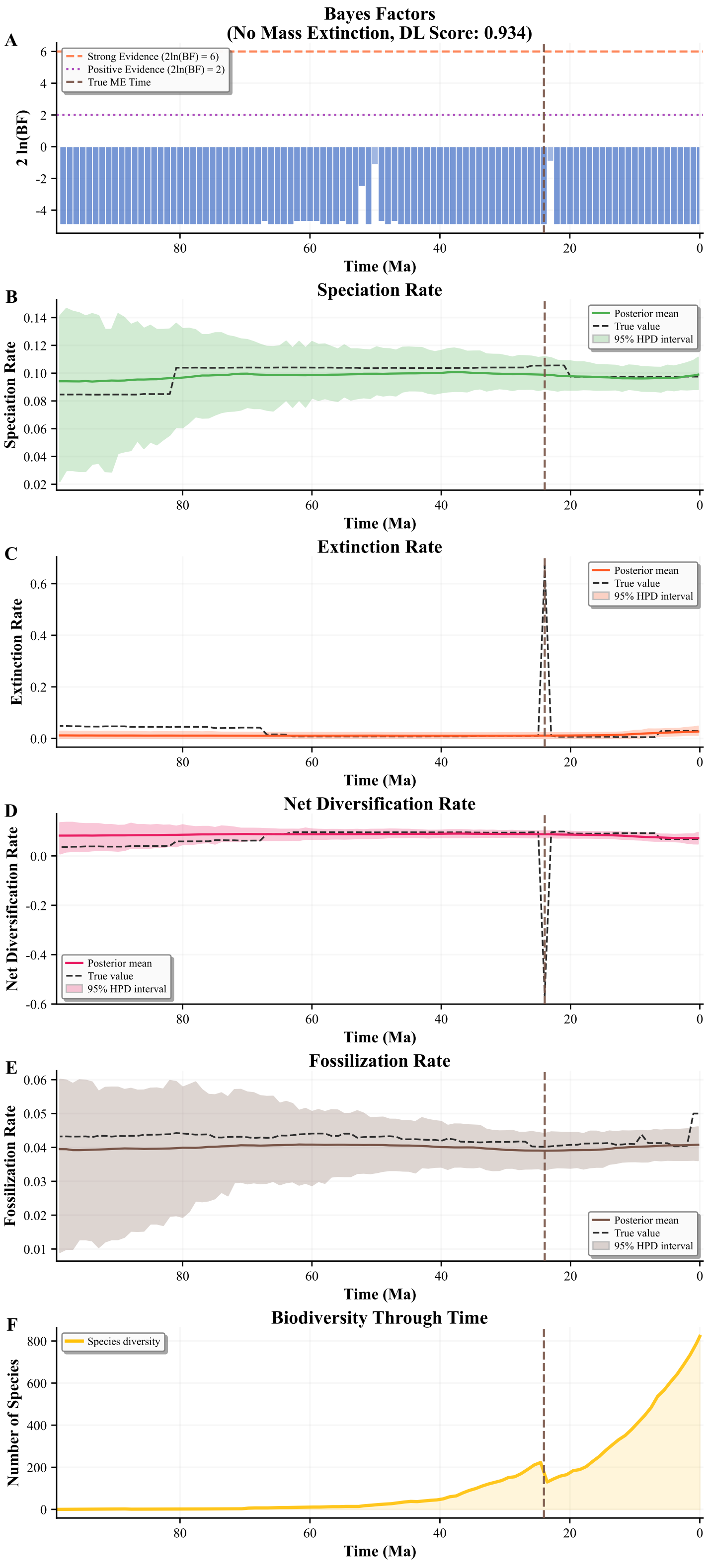


Figure S11: Deep learning and Bayesian inference for a simulated evolutionary history. (A) deep learning score and Bayes factor, (B-E) key evolutionary rates (speciation, extinction, diversification, and fossilization), and (F) the overall trajectory of species diversity through time.

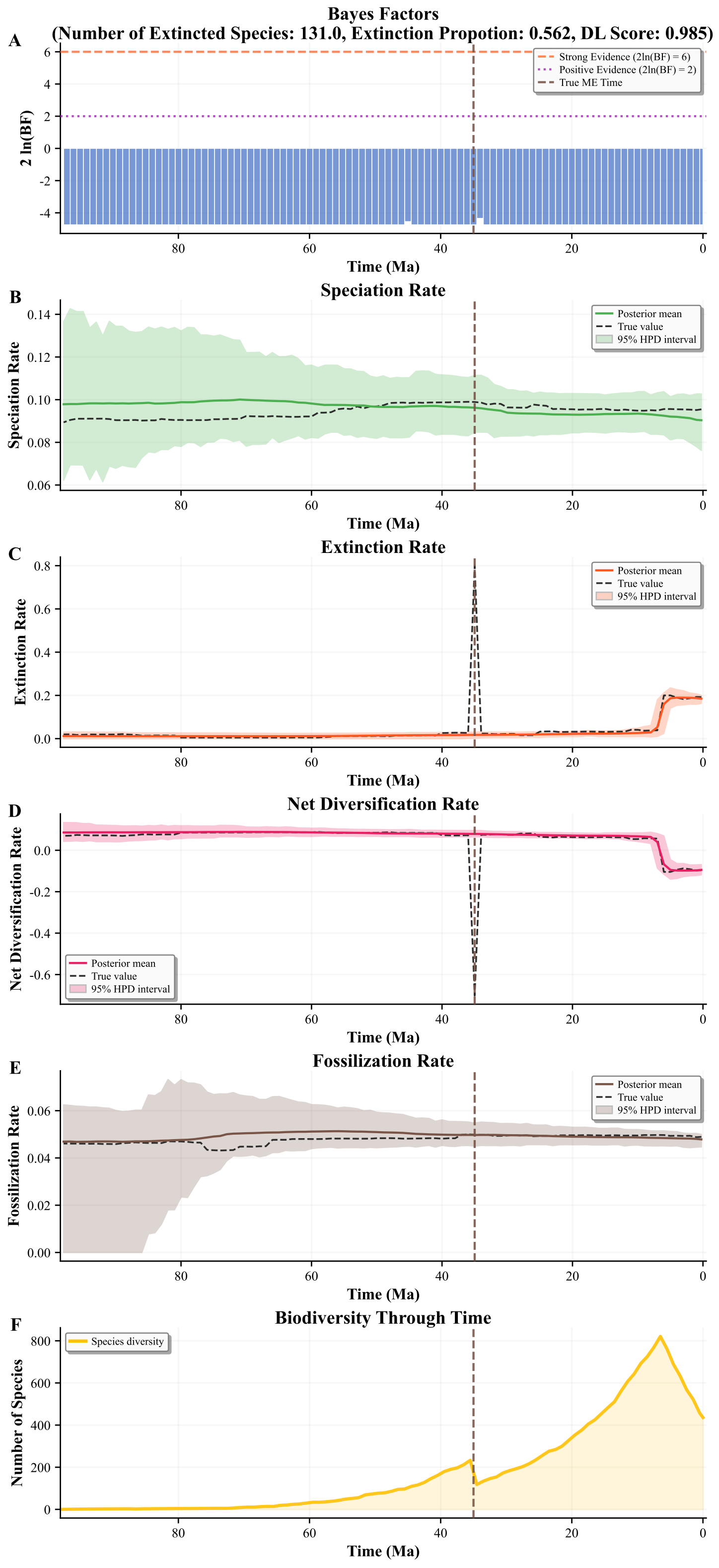


Figure S12: Deep learning and Bayesian inference for a simulated evolutionary history. (A) deep learning score and Bayes factor, (B-E) key evolutionary rates (speciation, extinction, diversification, and fossilization), and (F) the overall trajectory of species diversity through time.

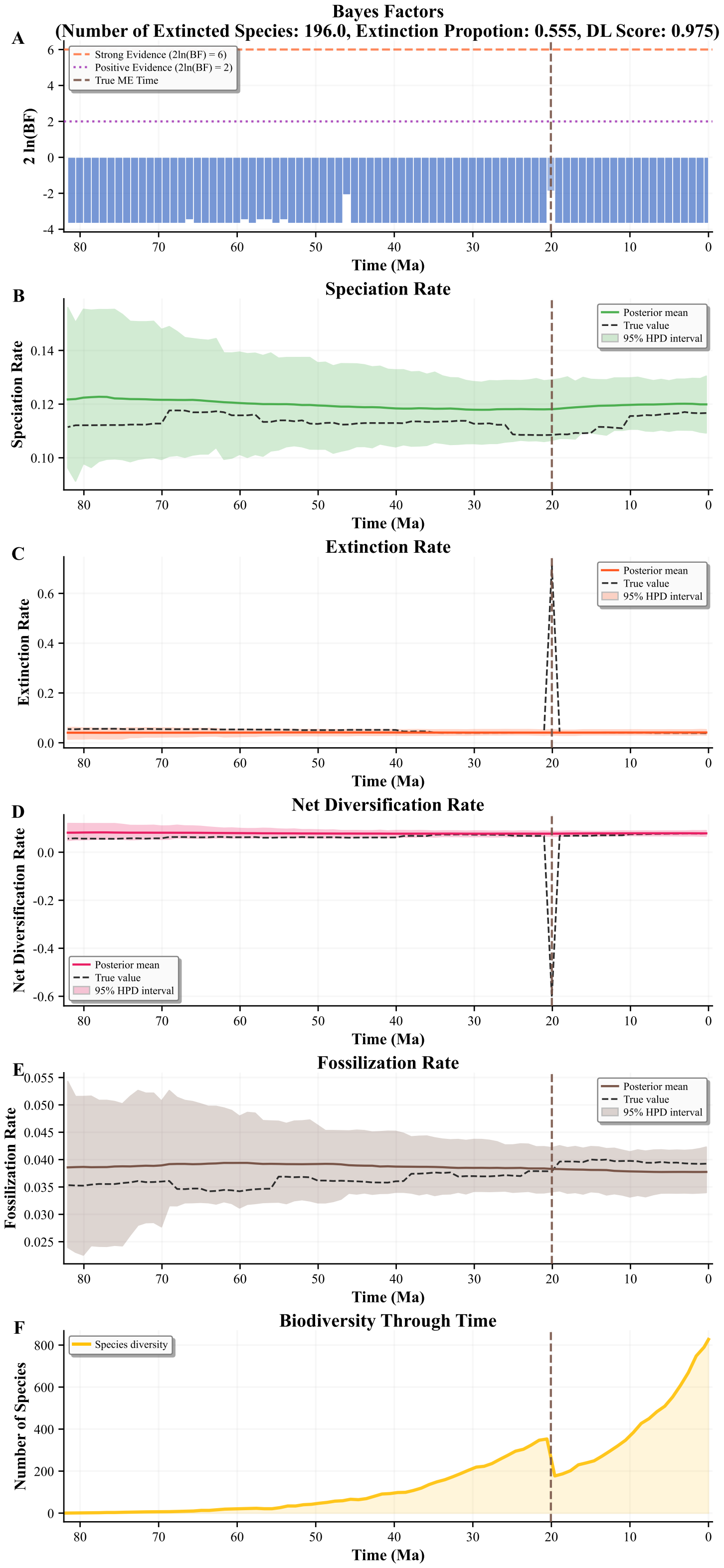


Figure S13: Deep learning and Bayesian inference for a simulated evolutionary history. (A) deep learning score and Bayes factor, (B-E) key evolutionary rates (speciation, extinction, diversification, and fossilization), and (F) the overall trajectory of species diversity through time.

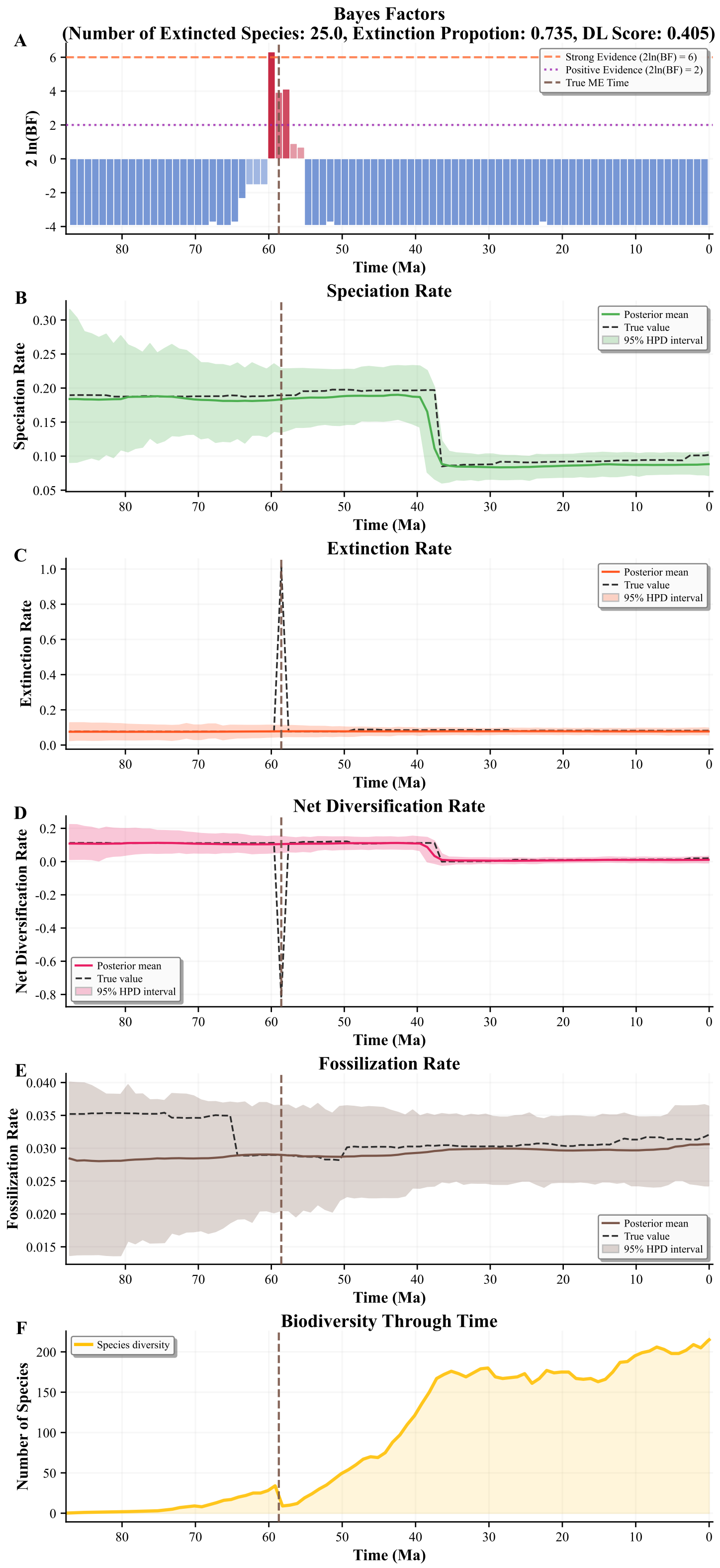


Figure S14: Deep learning and Bayesian inference for a simulated evolutionary history. (A) deep learning score and Bayes factor, (B-E) key evolutionary rates (speciation, extinction, diversification, and fossilization), and (F) the overall trajectory of species diversity through time.

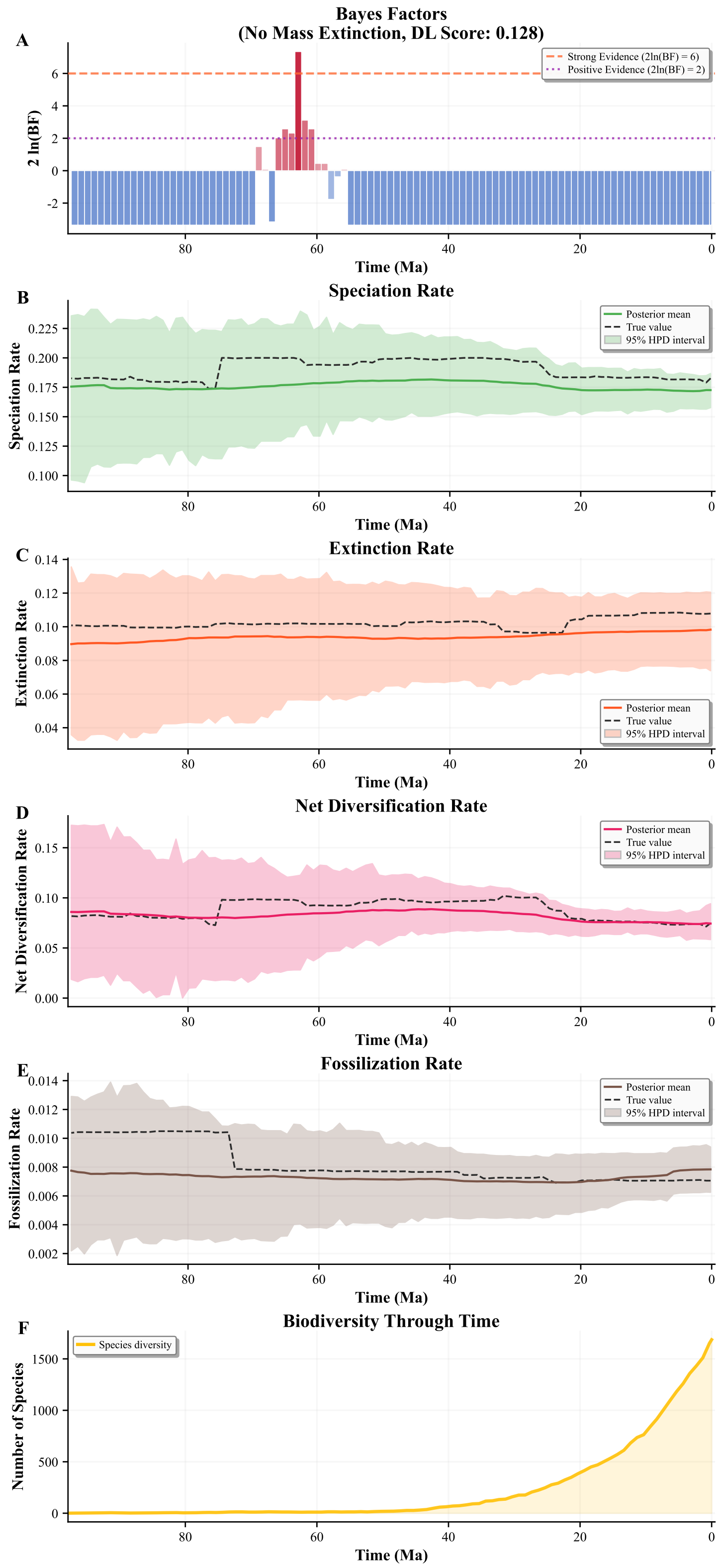


Figure S15: Deep learning and Bayesian inference for a simulated evolutionary history. (A) deep learning score and Bayes factor, (B-E) key evolutionary rates (speciation, extinction, diversification, and fossilization), and (F) the overall trajectory of species diversity through time.

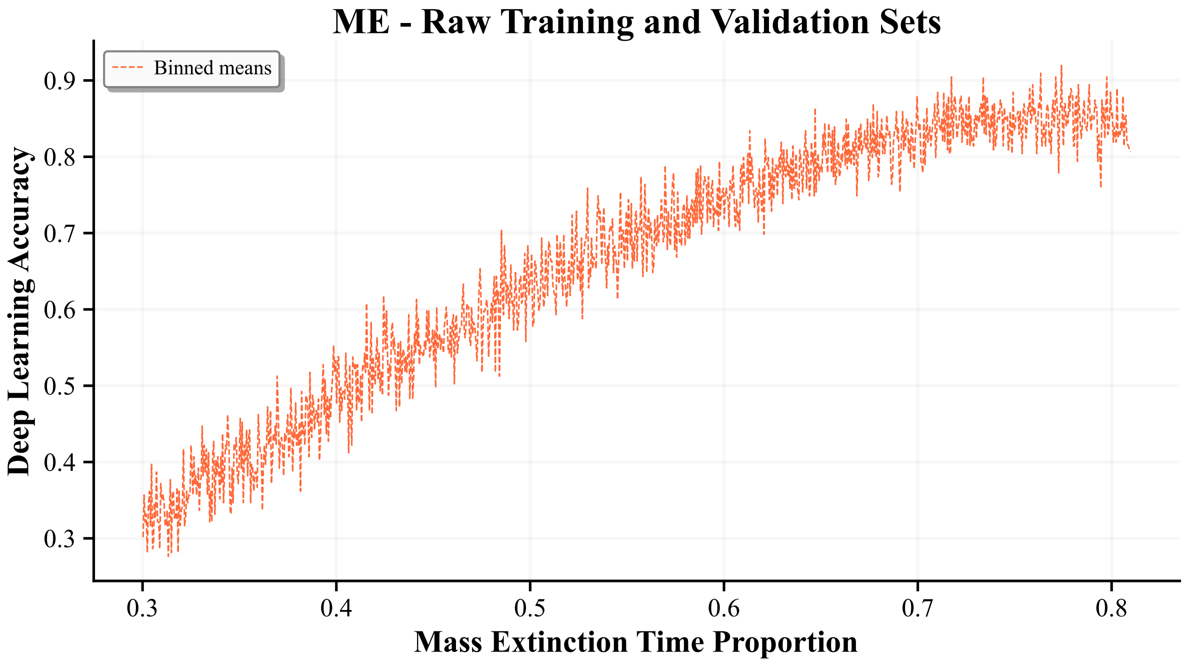


Figure S16: The impact of the relative time of mass extinction (time of mass extinction / origin time) on the predictive score and accuracy of the deep learning model. The performance trends are presented for scenarios with mass extinction (ME). The trend is evaluated and visualized for the training and validation datasets. The methods used to generate the binned means are detailed in Section S9. Note that mass extinction is determined here by the simulation itself, not by our later redefinition.

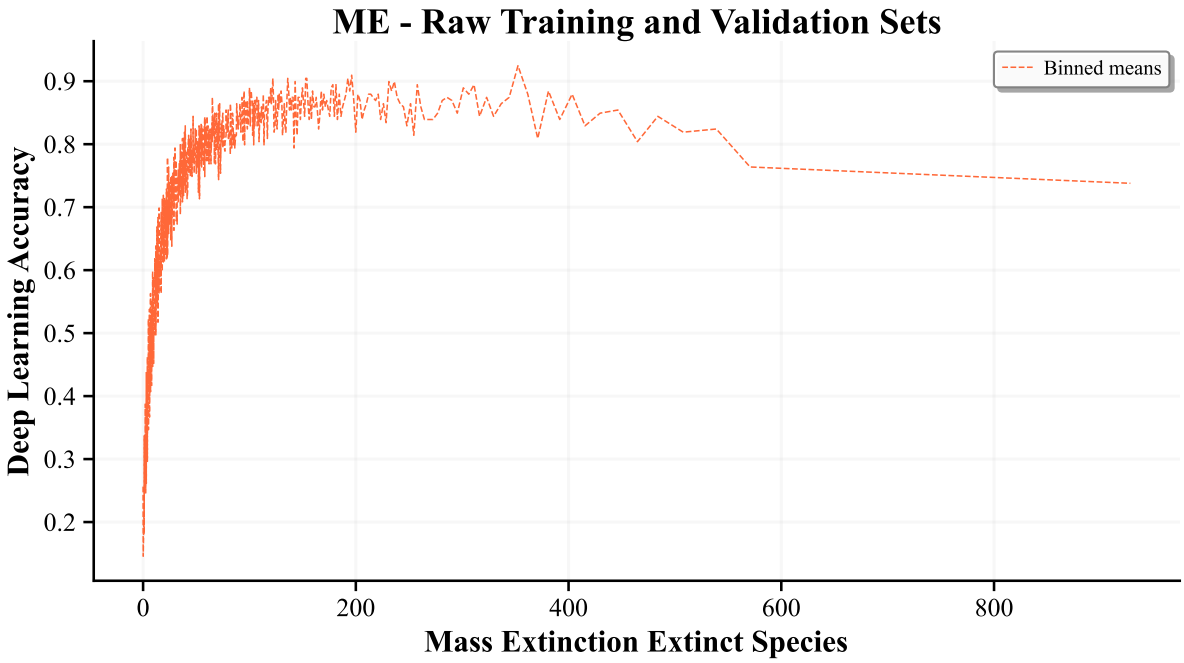


Figure S17: The impact of the number of extinct species of mass extinction on the predictive score and accuracy of the deep learning model. The performance trends are presented for scenarios with mass extinction (ME). The trend is evaluated and visualized for the training and validation datasets. The methods used to generate the binned means are detailed in Section S9. Note that mass extinction is determined here by the simulation itself, not by our later redefinition.

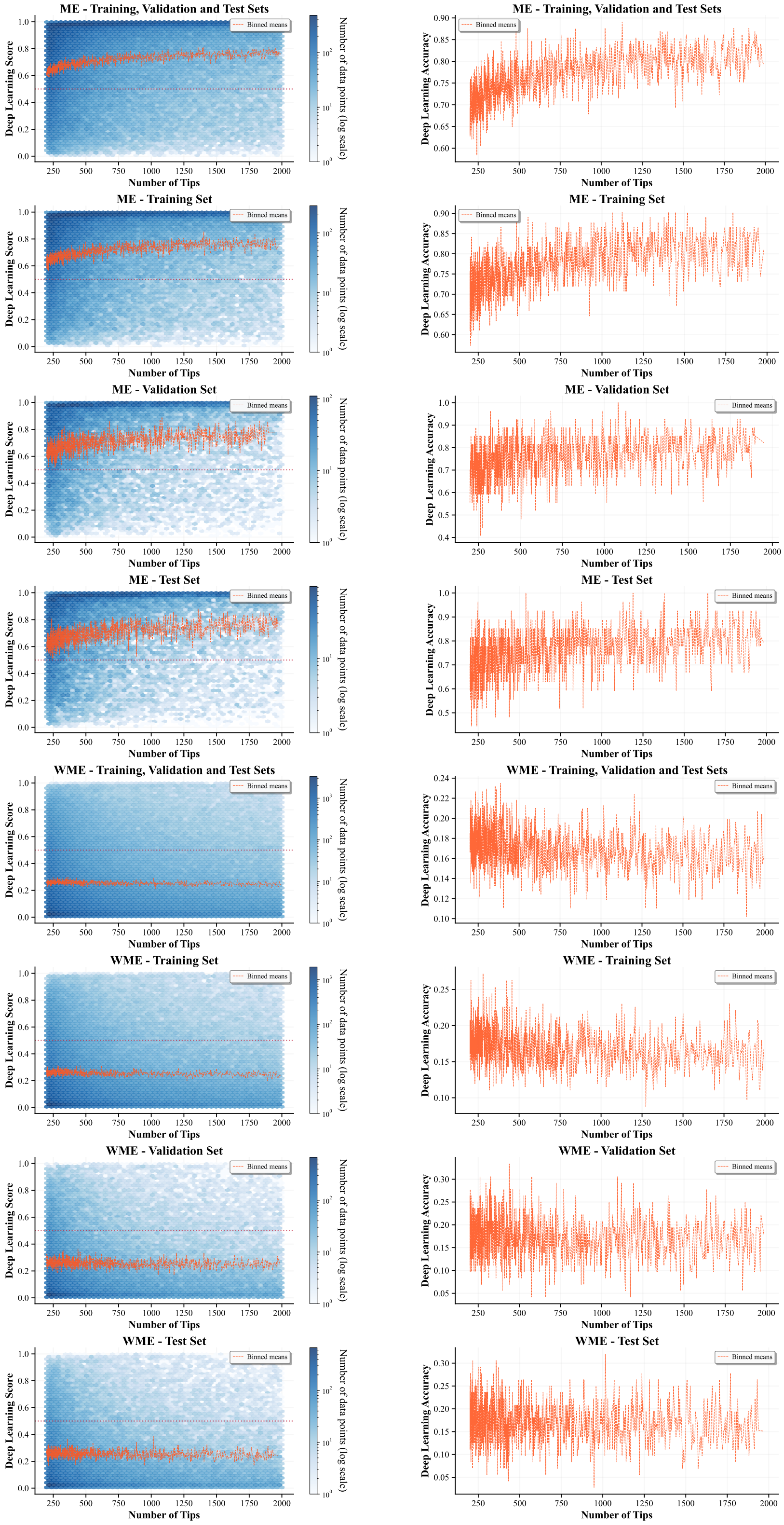


Figure S18: The impact of the number of tips on the predictive score and accuracy of the deep learning model. The performance trends are presented for scenarios both with mass extinction (ME) and without mass extinction (WME). Each trend is independently evaluated and visualized for the training, validation, and test datasets. The methods used to generate the binned means are detailed in Section S9.

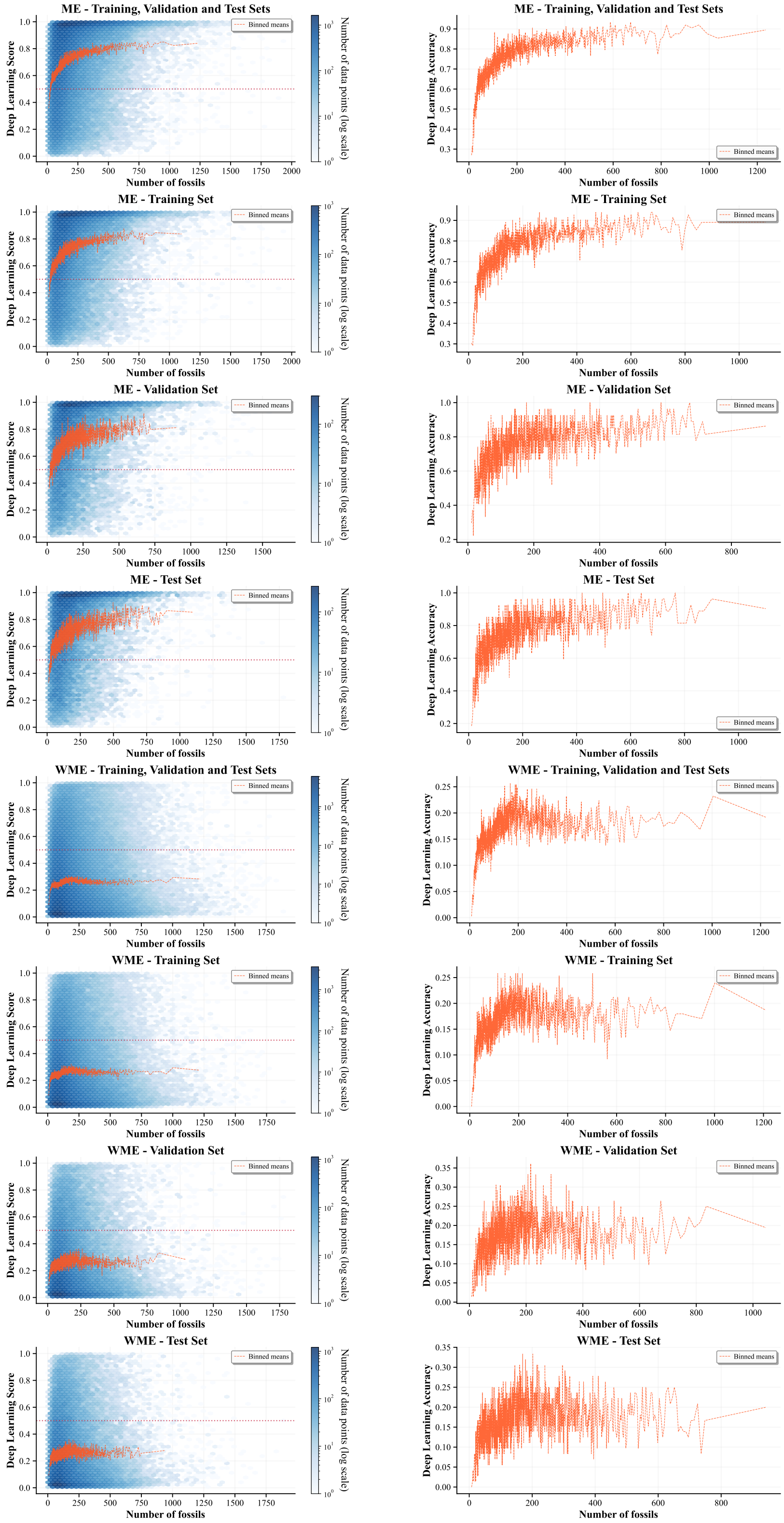


Figure S19: The impact of the number of fossils on the predictive score and accuracy of the deep learning model. The performance trends are presented for scenarios both with mass extinction (ME) and without mass extinction (WME). Each trend is independently evaluated and visualized for the training, validation, and test datasets. The methods used to generate the binned means are detailed in Section S9.

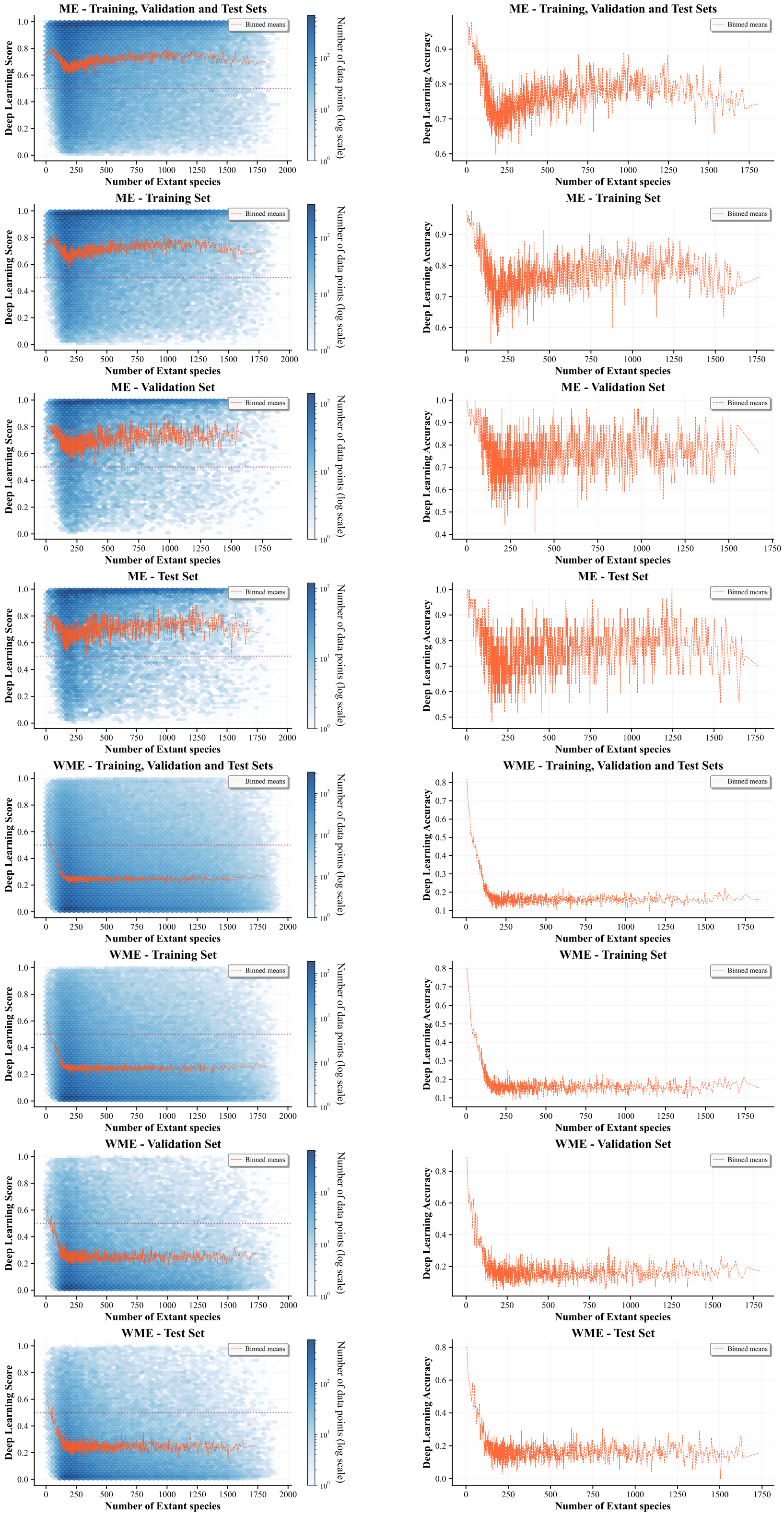


Figure S20: The impact of the number of extant species on the predictive score and accuracy of the deep learning model. The performance trends are presented for scenarios both with mass extinction (ME) and without mass extinction (WME). Each trend is independently evaluated and visualized for the training, validation, and test datasets. The methods used to generate the binned means are detailed in Section S9.

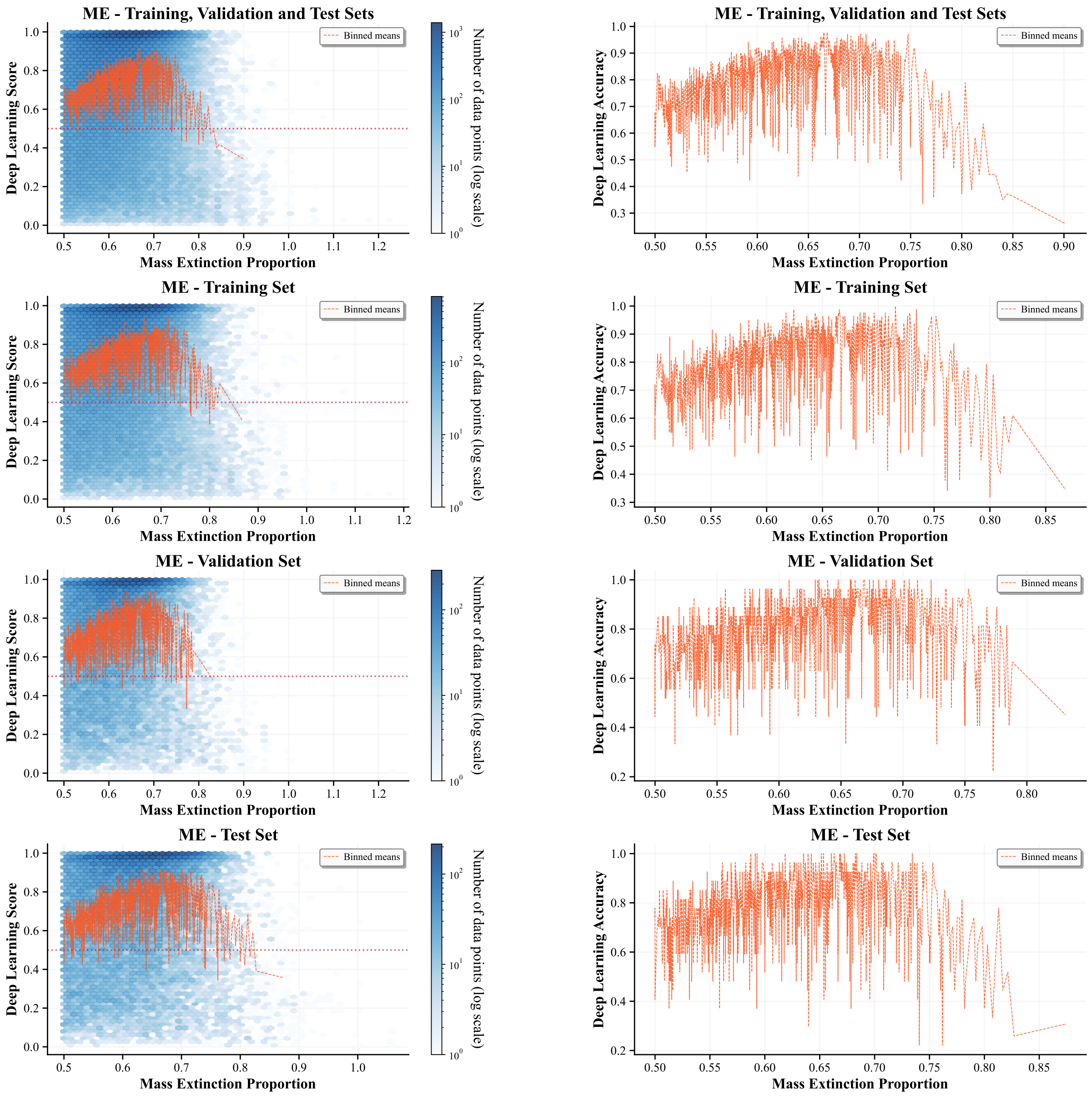


Figure S21: The impact of the extinction proportion of mass extinction on the predictive score and accuracy of the deep learning model. The performance trends are presented for scenarios both with mass extinction (ME) and without mass extinction (WME). Each trend is independently evaluated and visualized for the training, validation, and test datasets. The methods used to generate the binned means are detailed in Section S9.

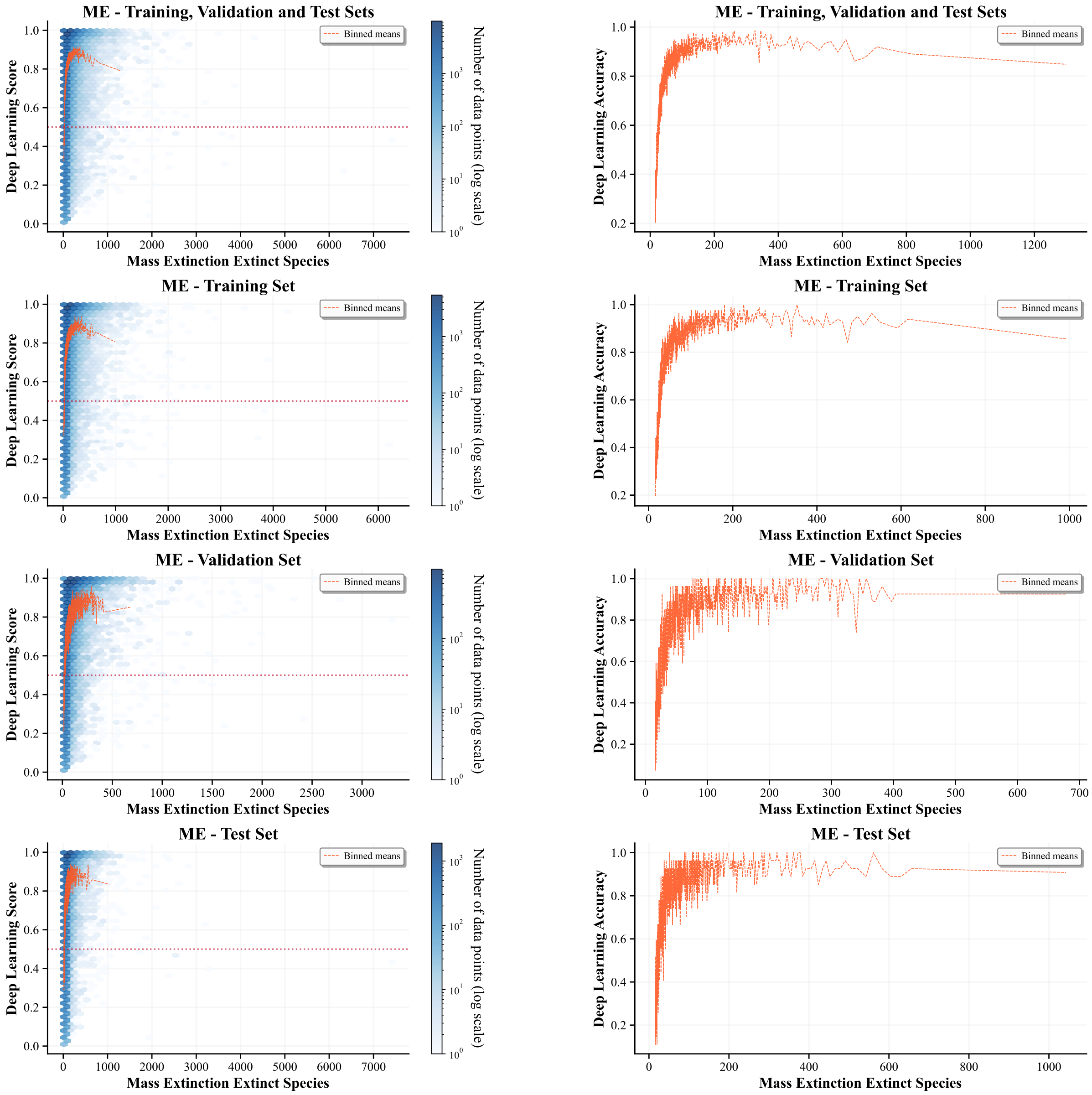


Figure S22: The impact of the number of extinct species of mass extinction on the predictive score and accuracy of the deep learning model. The performance trends are presented for scenarios both with mass extinction (ME) and without mass extinction (WME). Each trend is independently evaluated and visualized for the training, validation, and test datasets. The methods used to generate the binned means are detailed in Section S9.

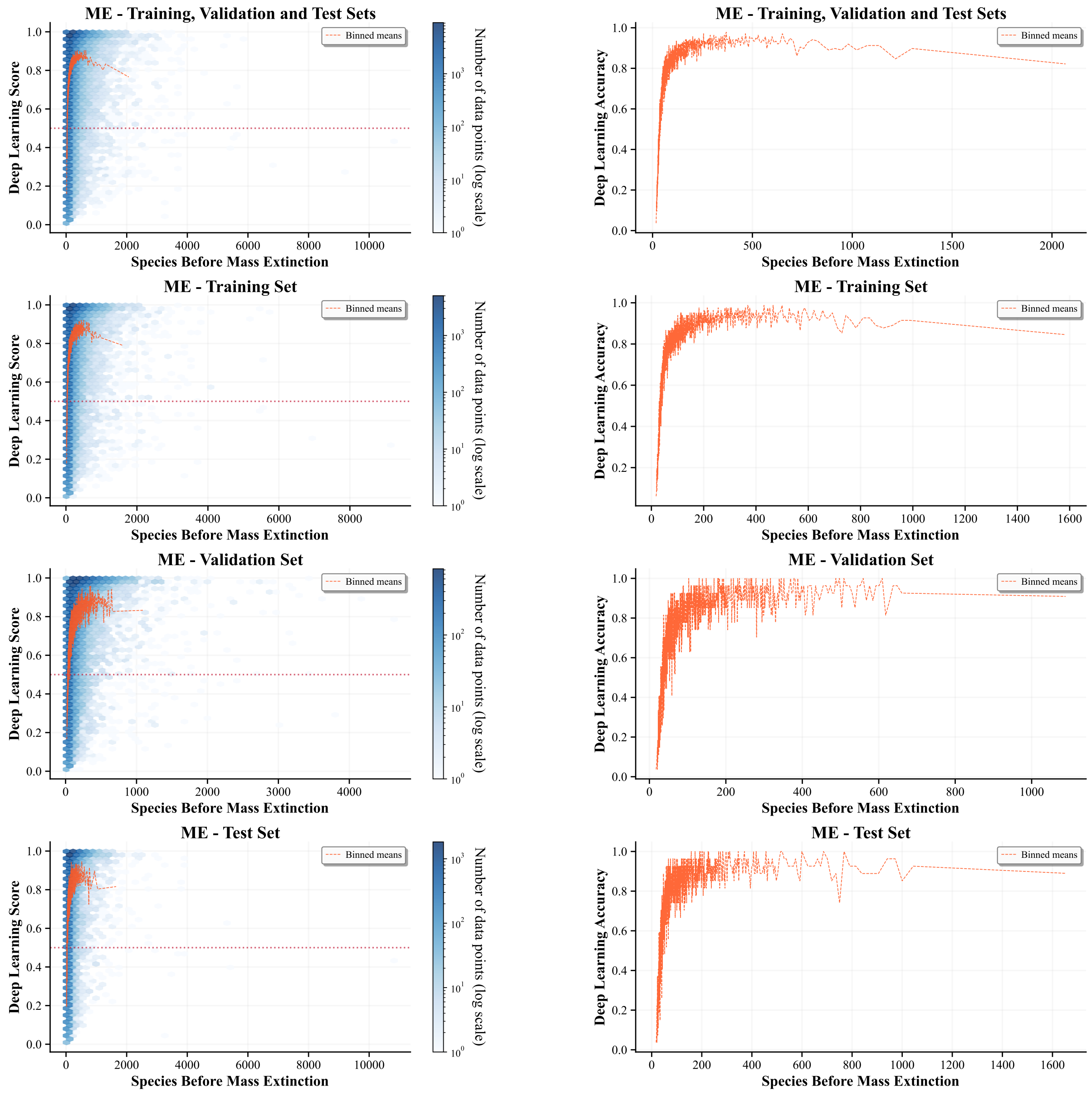


Figure S23: The impact of the number of species before mass extinction on the predictive score and accuracy of the deep learning model. The performance trends are presented for scenarios both with mass extinction (ME) and without mass extinction (WME). Each trend is independently evaluated and visualized for the training, validation, and test datasets. The methods used to generate the binned means are detailed in Section S9.

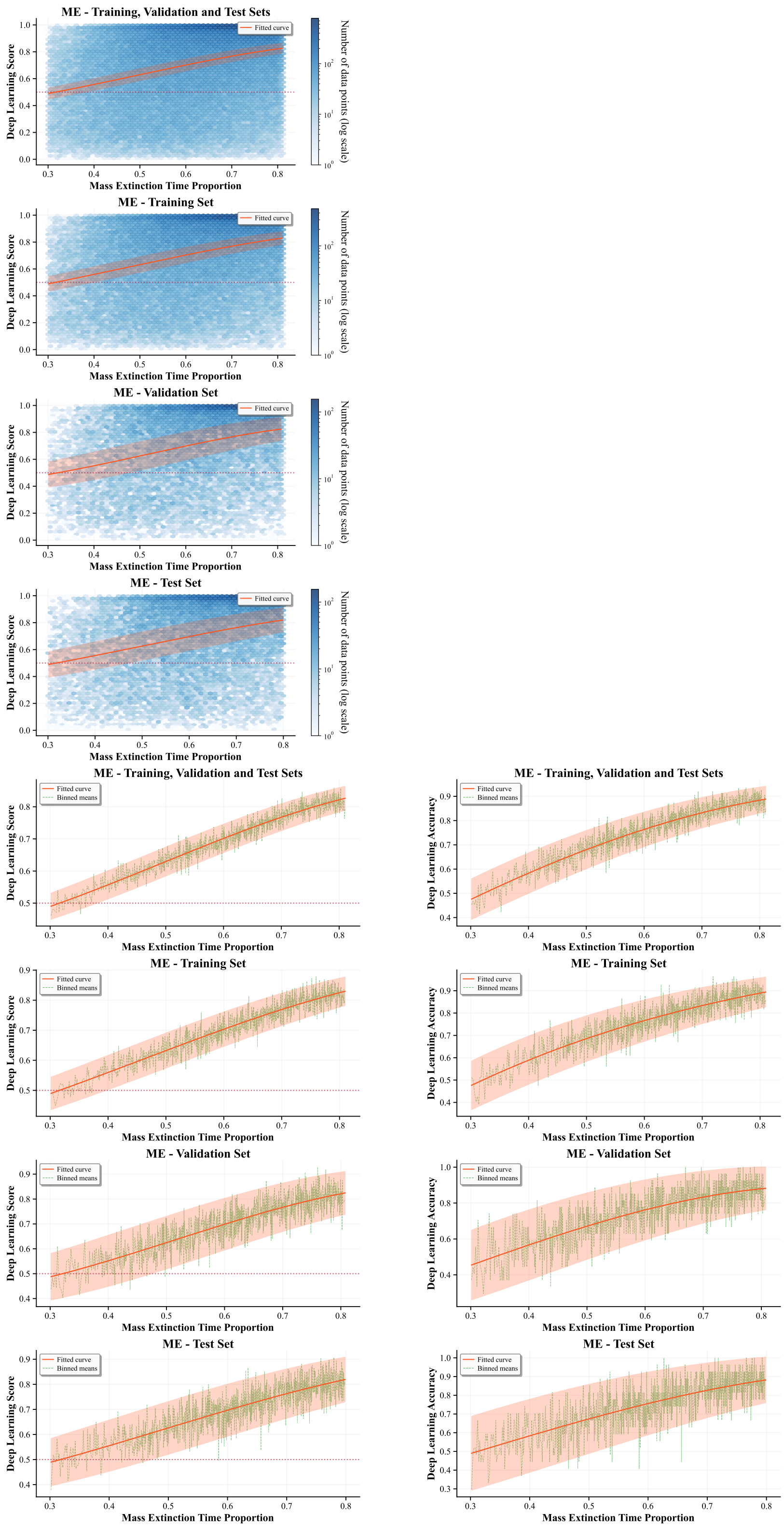


Figure S24: The impact of the relative time of mass extinction (time of mass extinction / origin time) on the predictive score and accuracy of the deep learning model. The performance trends are presented for scenarios both with mass extinction (ME) and without mass extinction (WME). Each trend is independently evaluated and visualized for the training, validation, and test datasets. The methods used to generate the fitted curves and their corresponding 95% confidence bands are detailed in Section S9.

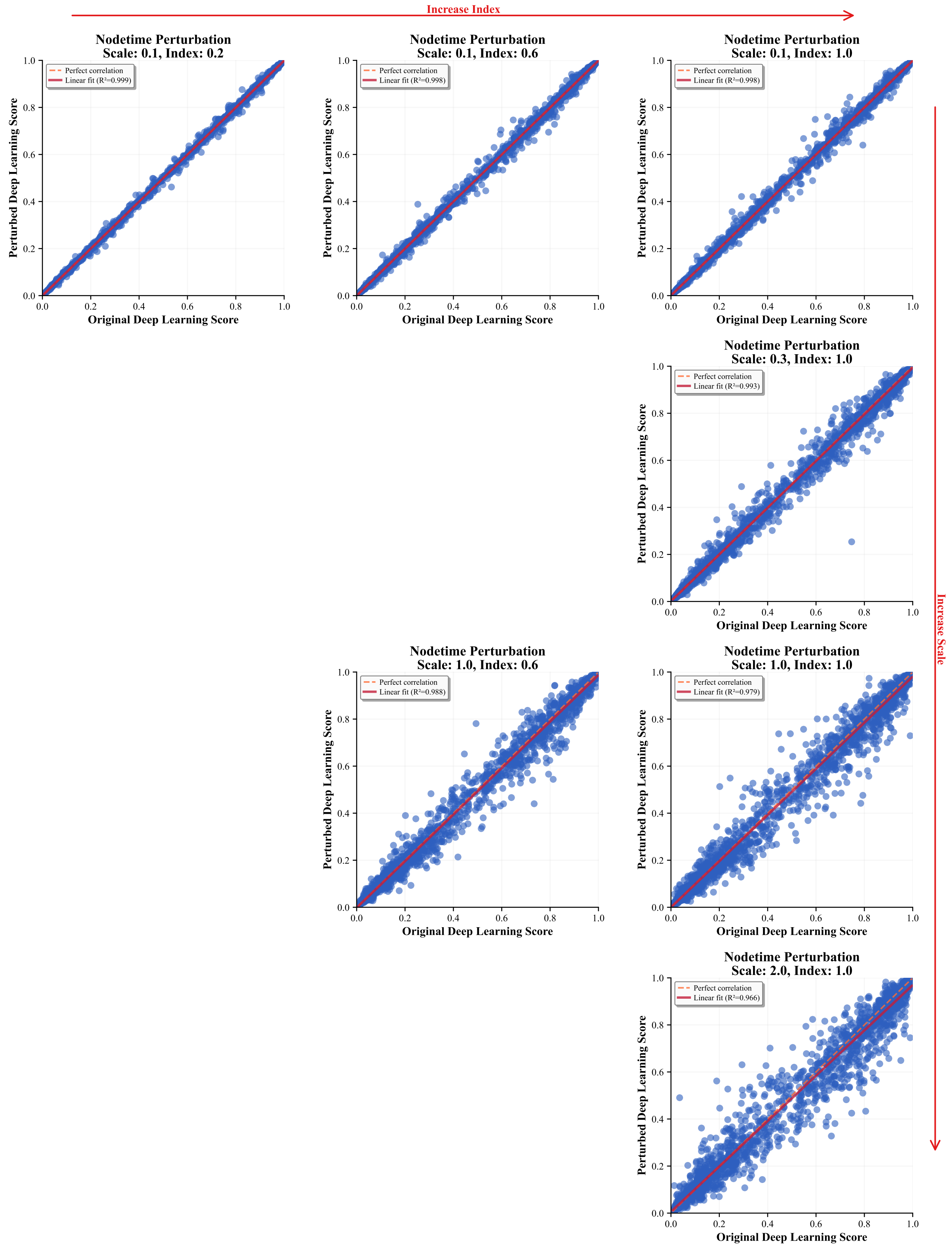


Figure S25: Robustness of the model's mass extinction score to perturbations of internal node ages. The analysis varies both the magnitude of the age shifts (controlled by the `scaleFactor`) and the proportion of internal nodes that are modified (the perturbation index). For a detailed description of the perturbation scheme, see Section S7.1.

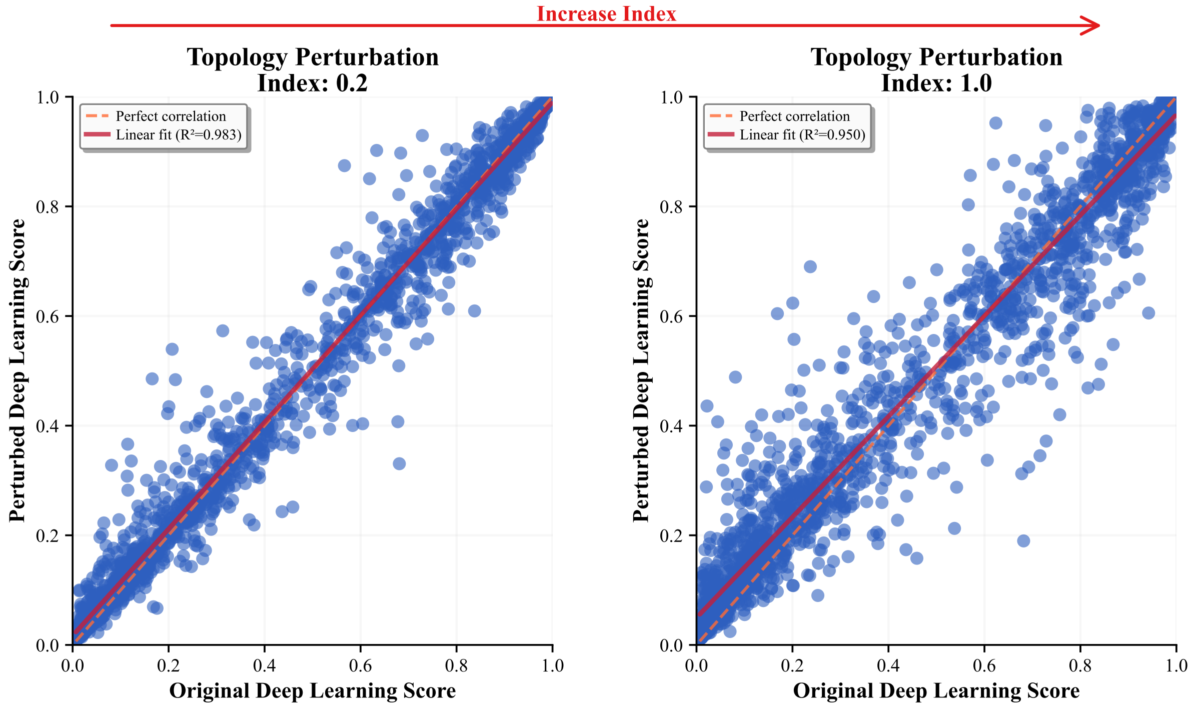


Figure S26: Robustness of the model's mass extinction score to perturbations of topology. The analysis varies the proportion of topology (the perturbation index). For a detailed description of the perturbation scheme, see Section S7.2.

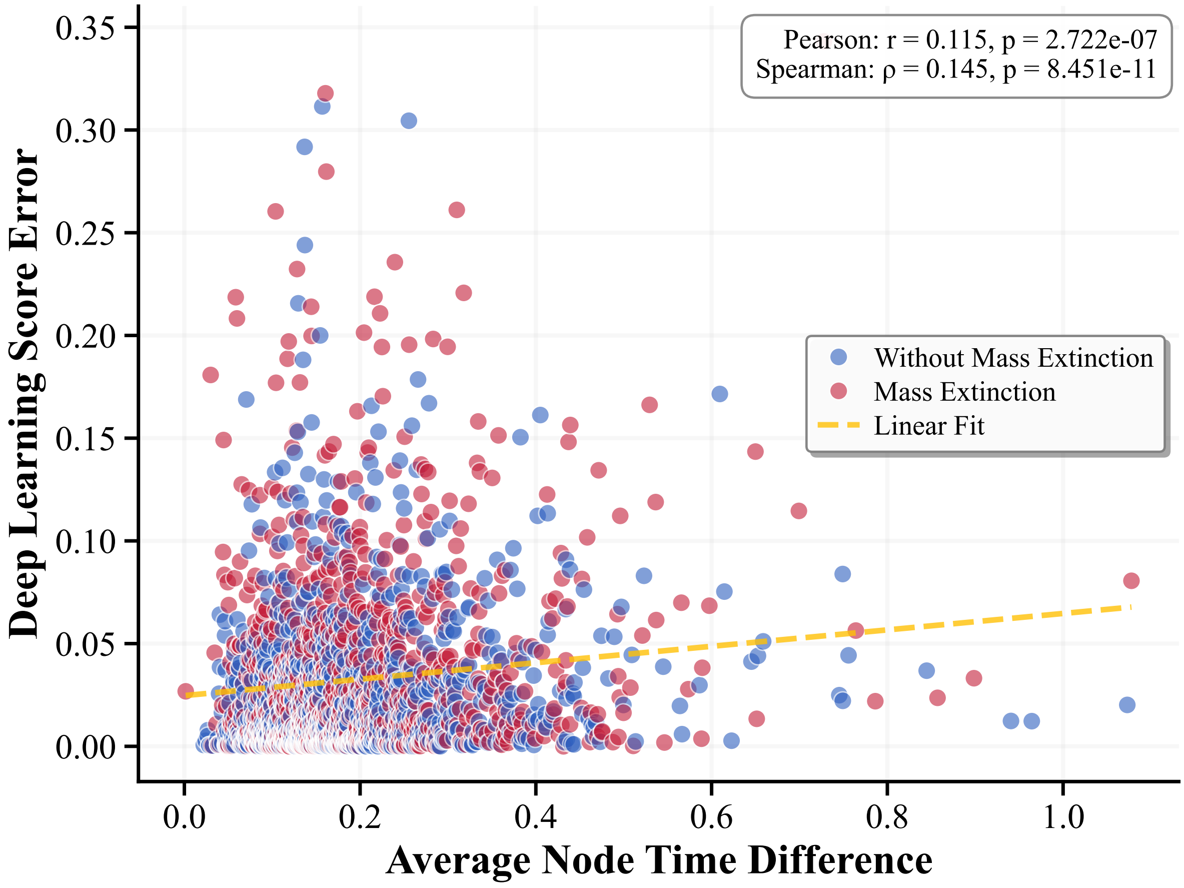


Figure S27: Scatter plot showing the relationship between the mean absolute change in internal node ages per tree and the corresponding change in the predicted mass extinction score.

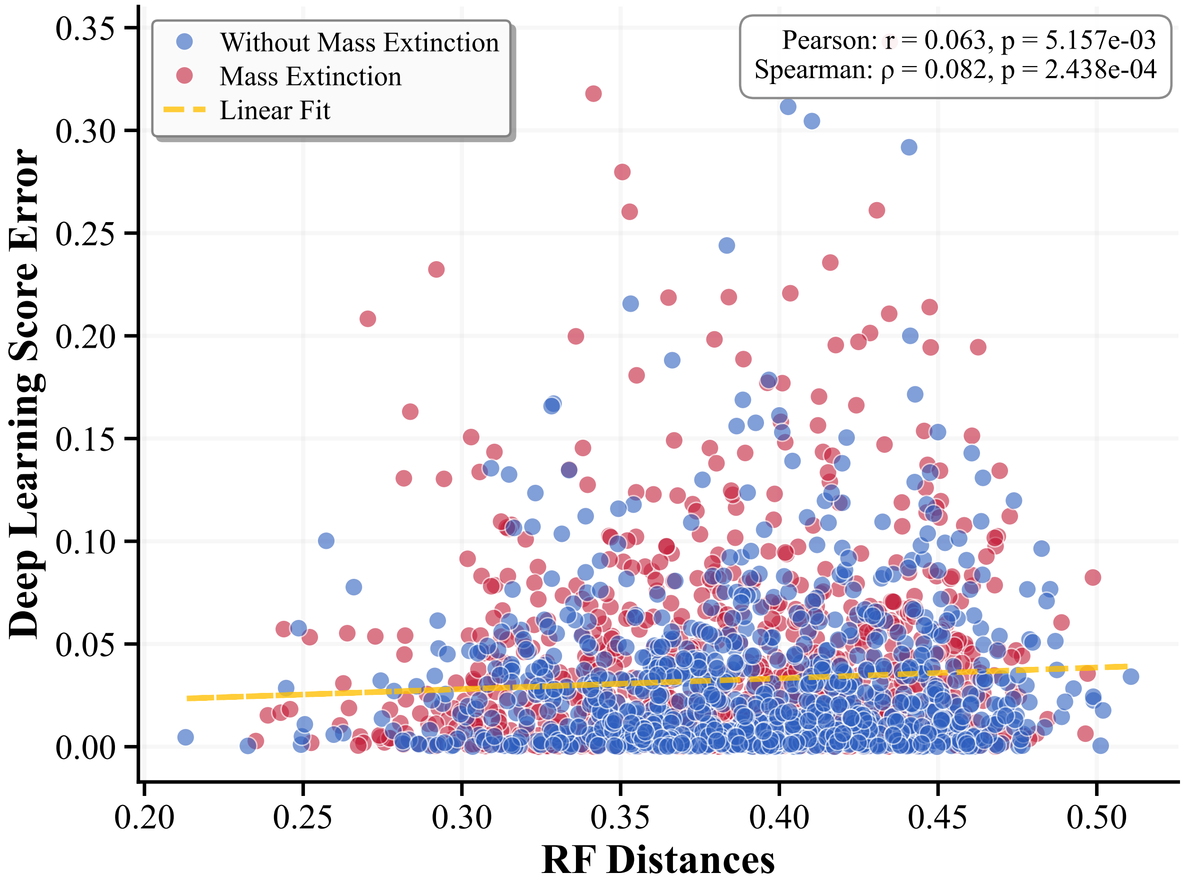


Figure S28: Scatter plot showing the relationship between the mean absolute change in RF distances per tree and the corresponding change in the predicted mass extinction score.
